# Widespread Peptide Surfactants with Post-translational *C-*methylations Promote Bacterial Development

**DOI:** 10.1101/2024.01.23.576971

**Authors:** Chen Zhang, Mohammad R. Seyedsayamdost

## Abstract

Bacteria produce a variety of peptides to mediate nutrient acquisition, microbial interactions, and other physiological processes. Of special interest are surface-active peptides that aid in growth and development. Herein, we report the structure and characterization of clavusporins, unusual and hydrophobic ribosomal peptides with multiple *C*-methylations at unactivated carbon centers, which help drastically reduce the surface tension of water and thereby aid in *Streptomyces* development. The peptides are synthesized by a previously uncharacterized protein superfamily, termed DUF5825, in conjunction with a vitamin B_12_-dependent radical *S*-adenosylmethionine metalloenzyme. The operon encoding clavusporin is wide-spread among actinomycete bacteria, suggesting a prevalent role for clavusporins as morphogens in erecting aerial hyphae and thereby advancing sporulation and proliferation.

## Main Text

Natural products are small organic molecules that are synthesized by a wide array of organisms for diverse purposes^1^. Their functions fall into several broad categories, many involving chemical warfare or otherwise competitive or collaborative associations^1–5^. An interesting class are those that interact with the environment in a physical manner. Siderophores, for example, mine iron from insoluble sources and deliver the metal to the host^6^, while osmolytes enhance the ionic strength of water^7^. Another intriguing class are surfactants, which can alter the surface tension of water and allow microbes to raise aerial structures, among other processes^8,9^. Surfactants are especially important for the development of microbes, notably in streptomycetes and filamentous fungi, which produce aerial hyphae^10–15^. These structures develop into spores that, upon dispersion and germination, allow the respective life cycles to start anew^4^. Sporulation is a highly complex and coordinated process and the small molecules underlying it remain underexplored. The recent accumulation of genome sequencing data suggests that many natural product groups remain to be discovered, including those involved in microbial development^16^.

SapB is a ribosomally synthesized and post-translationally modified peptide (RiPP) in the lanthipeptide family and the only well-known surface-active natural product in *Streptomyces* development^17–19^. It is absent or highly divergent in many actinomycete genomes, suggesting that alternative surfactants may be involved in this process. We became interested in the involvement of surfactant peptides in *Streptomyces clavuligerus*, the producer of clavulanic acid, as it encodes a score of uncharacterized biosynthetic gene clusters, notably a RiPP cluster, the expression of which requires a rare leucine tRNA. This tRNA is encoded by *bldA* and its presence is a marker for genes that have functions related to aerial hyphae formation and sporulation^10–13^. The RiPP cluster is found on the megaplasmid pSCL4 in *S. clavuligerus* and prior studies have shown that removal of this plasmid leads to a sporulation-defective phenotype; however, the underlying molecular basis has remained elusive^20^. We have herein investigated the product of this RiPP gene cluster and found it to be a novel family of peptides that are synthesized by a highly unusual mechanism. They are encoded in diverse actinomycete genomes and effectively reduce the surface tension water, thus acting as prevalent biosurfactants that advance microbial development.

### An unusual RiPP operon in *S. clavuligerus*

The *S. clavuligerus* genome encodes a RiPP gene cluster that we have termed *mpc* (**m**ethylated **p**eptides in *S. **c**lavuligerus*) (**Fig. 1a**). It contains a transporter (*mpcT*), an S9 peptidase (*mpcP*), a 39mer precursor peptide (*mpcA*), a vitamin B_12_-dependent radical *S-*adenosylmethionine (rSAM) enzyme (*mpcB*), and a domain of unknown function (DUF5825, *mpcC*). The cluster is notable for the presence of a rare TTA leucine codon in *mpcT*, which suggests an involvement in sporulation, as well as the rSAM enzyme MpcB, and the DUF5825 MpcC. The hypothetical protein MpcC has no sequence similarity with known proteins; members of DUF5825 have not yet been characterized. MpcC does show high structural similarity with the *C*-terminus of MpcB (at residues 356-612) by HHPred analysis, implying functional relevance between these two proteins (**Fig. S1**)^21^. rSAM enzymes utilize a [4Fe-4S] cluster to generate a highly reactive 5ʹ-deoxyadenosyl radical (5ʹ-dA•), a radical initiation process that underlies many unusual transformations in biology^22–24^. A subfamily of rSAM enzymes bind B_12_. These so-called class B rSAM methyltransferases can methylate unactivated *sp*^3^-carbons^25–27^. Several RiPPs have been reported with methyl groups installed by B_12_-rSAM enzymes, including polytheonamides, bottromycins and pheganomycin^28–32^. Unlike canonical rSAM enzymes, which share a conserved CX_3_CX_2_C motif for [4Fe-4S] cluster binding, MpcB contains a CX_7_CX_2_C binding sequence^22^.

**Fig. 1.**
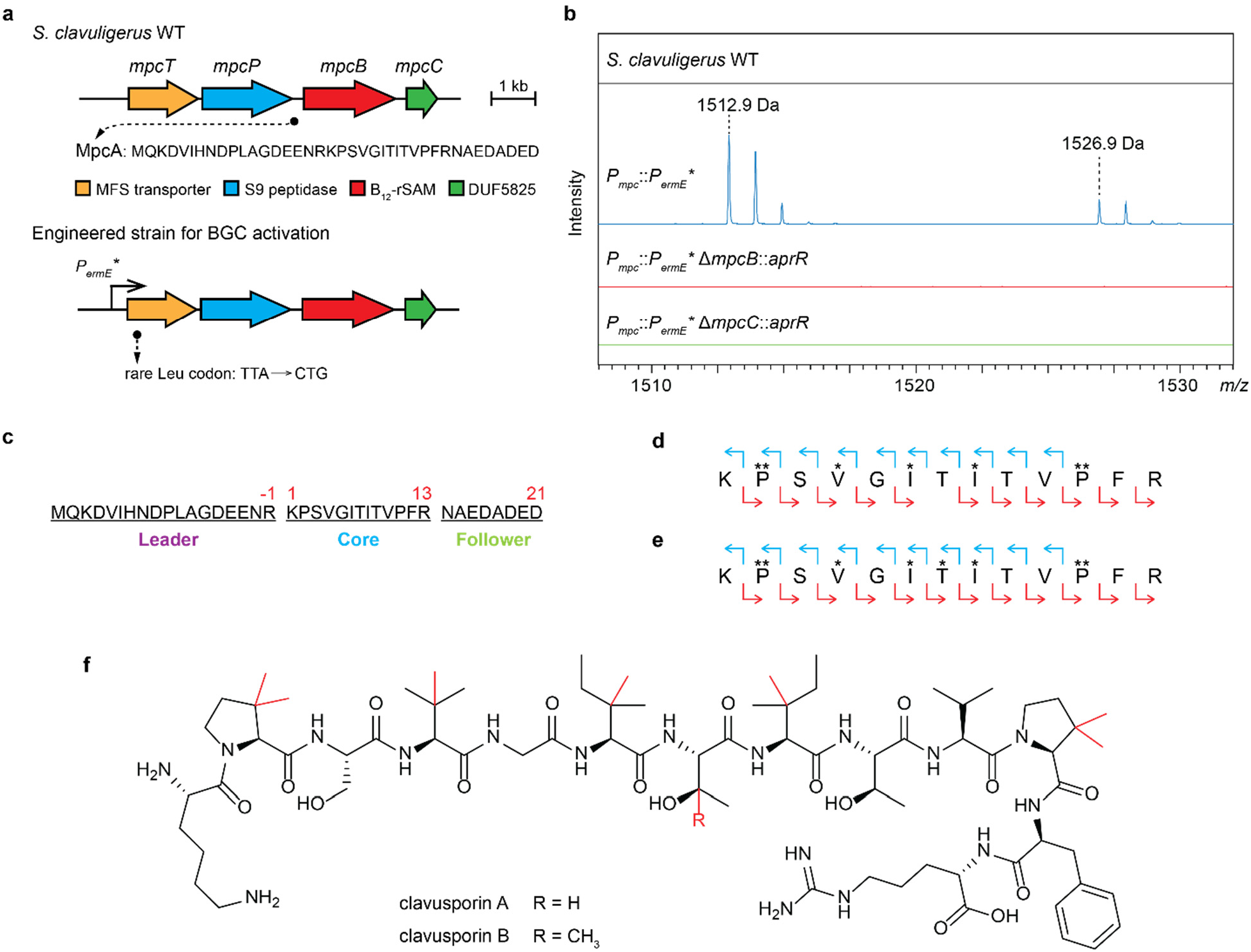
Discovery and structure of clavusporins. **a**, The *mpc* gene cluster as identified by genome mining. The gene locus of *mpcA* is marked by a black circle, and its protein sequence is shown. Strategies for genetic activation are shown below, including promoter exchange and codon optimization. **b**, MALDI-TOF MS analysis of SDS extracts of *S. clavuligerus* strains indicated. **c**, Numbering and parts of the MpcA peptide. **d-e**, MS/MS profiles of clavusporin A (**d**) and B (**e**). Blue and red arrows represent observed b and y ions, respectively. Each asterisk represents methylation on the corresponding residue. **f**, Structure of clavusporins. Post-translational methylations are marked in red.

To circumvent any transcriptional regulation of the *mpc* operon and identify its product, we engineered an *S. clavuligerus* strain, wherein the native promoter *P_mpc_* was replaced with a *Streptomyces* constitutive promoter *P_ermE_\** and the first 33 basepairs of *mpcT* were codon-optimized, including a switch from the rare Leu codon (TTA) to a common one (CAG) (**Fig. 1a**, **Fig. S2, Table S1**, see *Methods*)^33^. However, we were unable to identify methylated peptides from culture supernatants or organic extract of cell pellets of the engineered strain via untargeted peptidomic analysis. We reasoned that the potential methylations catalyzed by MpcB could drastically elevate the hydrophobicity of the peptide product(s), which could become recalcitrant to solvent extraction. Inspired by the discovery of SapB, the cell pellet of *S. clavuligerus P_mpc_::P_ermE_\** was extracted with 1% sodium dodecyl sulfate solution, and the extracts analyzed by matrix-assisted laser desorption/ionization time-of-flight (MALDI-TOF) mass spectrometry^34^. Two signals (*m/z* 1512.9 and 1526.9) were observed exclusively in the engineered strain but not WT (**Fig. 1b**). The 14 Da difference pointed to an additional methylation, further confirming the hypothesis that the gene cluster produces methylated peptides. Genetic deletion of either *mpcB* or *mpcC* in *S. clavuligerus P_mpc_*::*P_ermE_\** abolished the production of these 2 signals (**Fig. 1b** and **Fig. S3**), thus linking their production to the *mpc* locus. The two products match 13mer MpcA peptides (KPSVGITITVPFR, **Fig. 1c**) with 7 or 8 methylations; we have named these clavusporin A and B, respectively. Tandem HR-MS analysis revealed the methylation patterns (**Fig. 1d-e**), consisting of bismethylated Pro2 and Pro11, as well as monomethylated Val4, Ile6, and Ile8 on both peptides. Clavusporin B was additionally monomethylated at Thr7.

Extraction of the peptides into acidic ethyl acetate facilitated analysis by high-resolution mass spectrometry (HR-MS), the results of which were consistent with the assignment of hypermethylated 13mer peptides above (**Extended Fig. 1a-b**). To unequivocally determine the site of methylation, we isolated the clavusporins using a combination of unusual chromatographic methods and subjected them to multi-dimensional NMR spectral analysis, which revealed β-methylation at each modified residue, thus completing structural elucidation of these molecules (**Fig. 1f, Extended Fig. 1c-d**, **Table S2**, and **Fig. S4**). Clavusporins are a new family of β-methylated amphiphilic peptides; the occurrence of a series of β-methylated and β-dimethylated residues is rare in RiPP natural products^19,29,35^.

### Surface-active clavusporins promote bacterial development

The structures of clavusporins are unique in that the hydrophobic residues are heavily β-methylated, forming a tertiary alcohol on Thr7 and quaternary carbon centers on Pro, Val and Ile residues, while the side chains of Lys and Arg are positively charged under physiological conditions. We hypothesized that clavusporins can function as potent cationic surfactants in the course of *Streptomyces* development. To examine this hypothesis, we first measured the ability of clavusporins to reduce the surface tension of water using a contact angle goniometer. At a concentration of 1 μg/μL, clavusporin significantly altered the shape of aqueous droplets by reducing the surface tension to 31 mN/m, similar to the effect of soap (sodium dodecyl sulfate) at the same concentration (**Fig. 2a-c**)^36^. At lower concentrations, water surface tension decreased linearly with clavusporin titers (**Fig. 2c**, red line). Between 1.0–1.5 μg/μL, no further decreases were observed, indicating a critical micelle concentration of clavusporins at around 1 μg/μL. To test the effects of post-translational methylations, the unmodified 13mer core peptide, MpcA_1-13_, was synthesized (*Methods*, **Fig. S5** and **Table S3**) and subjected to surface tension analysis (**Fig. 2c**, blue line). At 1 μg/μL, MpcA_1-13_ only lowered surface tension to 57 mN/m, thus acting as a weaker surfactant and indicating that the methylations significantly affect the surfactant properties of clavusporin.

**Fig. 2.**
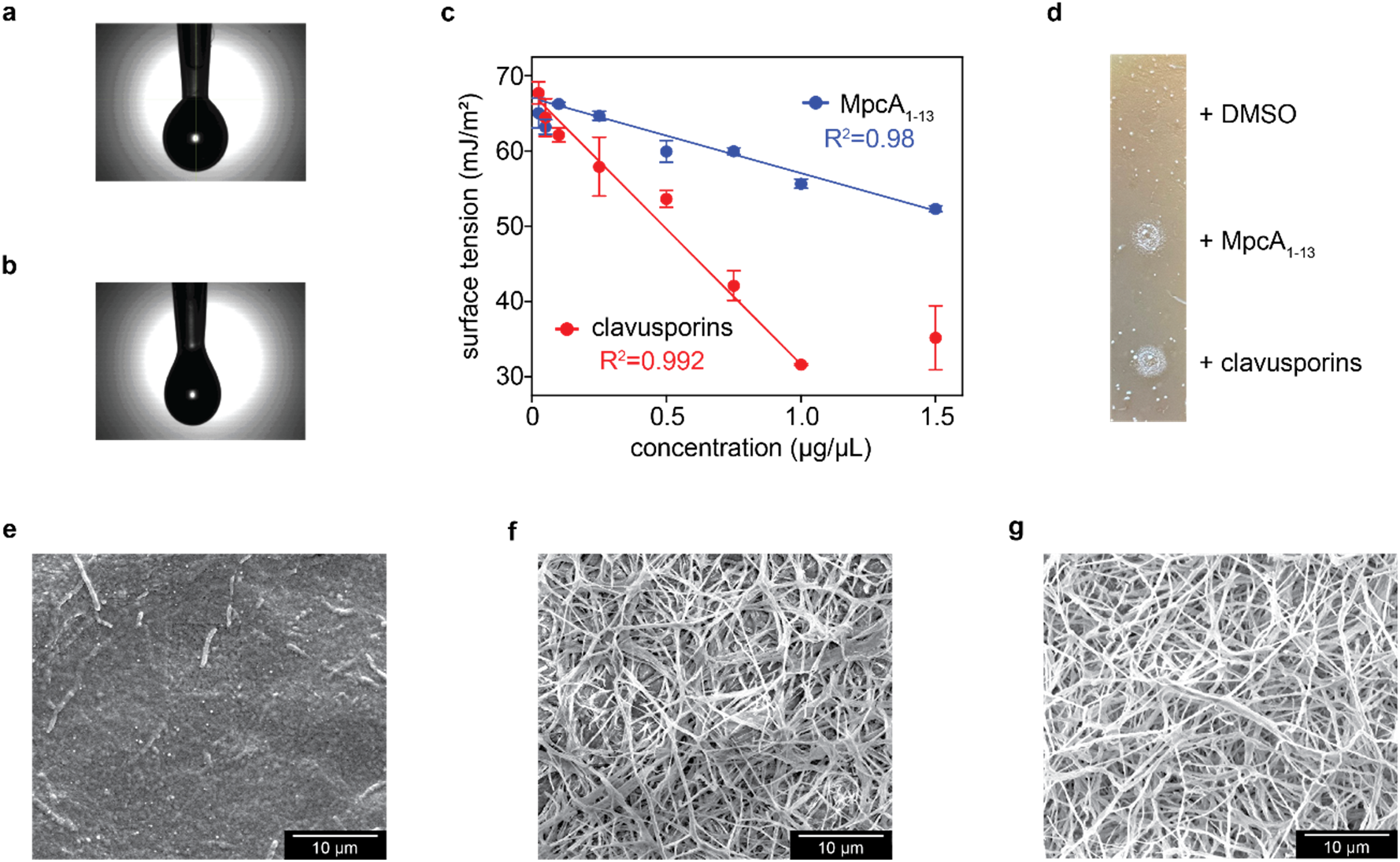
Clavusporins reduce surface tension and promote aerial hyphae growth. **a-b**, Images of a water droplet consisting of 10% DMSO (**a**, vehicle control) and a droplet of clavusporin at 1 μg/μL in a 10% DMSO solution (**b**). **c**, Quantification of surface-tension reduction by clavusporins (red) and the unmethylated 13mer, MpcA_1-13_ (blue). Surface tension is linearly dependent on compound concentration. The lowest surface tension occurs at ∼1 μg/μL clavusporins. **d**, Chemical complementation of *S. clavuligerus ΔmpcBC* with DMSO control, MpcA_1-13_, or clavusporins (10 μg dissolved in 2 μL of DMSO). **e-g**, SEM images of cell surfaces supplemented with DMSO control (**e**), MpcA_1-13_ (**f**), and clavusporins (**g**). Each experiment was repeated at least in three biological replicates, and representative results are shown.

We next assessed the effect of clavusporin on sporulation by generating gene inactivation mutants via replacement of *mpcBC* with an apramycin resistance cassette (*apr^R^*) (**Fig. S6**). The mutant was unable to sporulate, growing only substrate mycelia and limited aerial hyphae on agar, whereas WT cells developed aerial hyphae and spores under the same conditions (**Extended Fig. 2a-c**). These results are in line with prior reports of the developmental deficiency of *S. clavuligerus* mutants lacking the megaplasmid pSCL4^20^. The phenotype differences between WT and *ΔmpcBC* were verified by scanning electron microscopy (SEM) (**Extended Fig. 2d-f**). The WT showed ‘club’-like spore clusters, for which the strain has been named^37^; these were lacking in the mutant. These observations suggest that the *mpc* operon is involved in *S. clavuligerus* development through production of methylated peptides, consistent with the requirement of the rare leucine tRNA, encoded by *bldA*, for translation of MpcT.

To further test the function of clavusporin, we examined its ability to restore the bald phenotype of the *ΔmpcBC* strain. Cells growing on agar surfaces were either chemically supplemented with DMSO (as vehicle control) or with clavusporin. The clavusporin-treated area showed accelerated growth of aerial hyphae, based on visual inspection and SEM imaging (**Fig. 2d-g**), whereas the control did not. Moreover, complementation was dependent on clavusporin concentration, reaching levels of aerial hyphae growth that matched those of WT at 2 μg/μL clavusporin (Fig. S7).

Lastly, we explored the generality of these results by exploring the role of the orthologous *Streptomyces ghanaensis* (named *mgp* for methylated peptides in *S. ghanaensis*). This cluster encodes the same complement of genes with MpgA, MpgB, MpgC, MpgP, and MpgT proteins that are 77%, 81%, 58%, 63% and 56% homologous to those in *S. clavuligerus*, respectively (**Extended Fig. 3a**). We generated a *mpgB::apr^R^* insertion in *S. ghanaensis* (**Table S4**), which exhibited a white phenotype and altered development by visual inspection and by SEM. The WT, however, developed normally under similar conditions, producing dark grey spores (**Extended Fig. 3b-e** and **Fig. S8**). Together, these results show that the *mpc* operon produces peptide surfactants, which lower the surface tension of water and facilitate erection of aerial hyphae, thus playing a key role in the developmental processes of *S. clavuligerus* and *S. ghanaensis*. Whether they also play additional (auto)regulatory roles remains to be determined.

### Biosynthesis of clavusporins

With the structure and function of clavusporins established, we next explored its biosynthesis, focusing on the introduction of methyl groups at unactivated carbon centers. B_12_-dependent rSAM enzymes are notoriously difficult to purify, despite advances in cofactor incorporation, protein yields, and solubility of this class of enzymes^38,39^. We therefore utilized an alternative *in vivo* approach to co-express maltose-binding protein (MBP)-tagged MpcA with MpcB in *E. coli* cells (**Fig. 3a**, **Table S5**). Plasmids pDB1282 encoding Fe-S cluster assembly proteins and pBAD42-BtuCEDFB to enhance cellular cobalamin availability were also introduced into *E. coli*^39,40^. After expression, the MBP-tagged peptides were enriched through affinity-based chromatography and subsequently trimmed by proteolysis and analyzed by HPLC-coupled HR-MS.

**Fig. 3.**
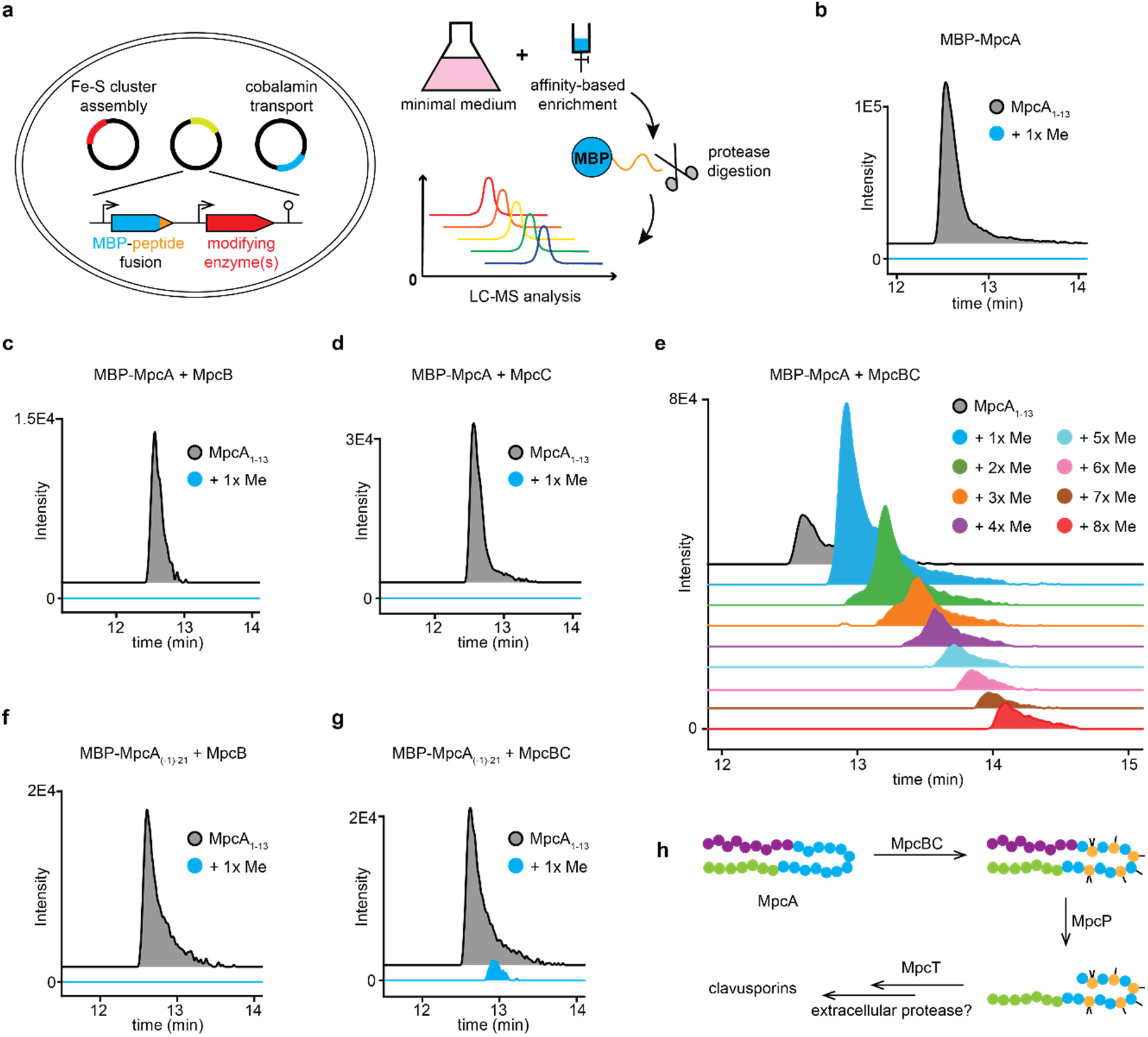
Biosynthesis of clavusporins. **a**, Workflow for *in vivo* reconstitution of peptide methylations by co-expression of MBP-tagged precursor peptide and the modification enzymes. **b-g**, HPLC-coupled HR-MS analysis of MpcA core peptides from MBP-peptide fusions after column-enrichment and trypsin digestion. Shown are extracted ion chromatograms of MpcA_1-13_ (core peptide) with or without methylation, as indicated. The traces are offset on the y axis for clarity. **h**, Proposed biosynthetic pathway of clavusporins.

As expected, only the linear, unmodified MpcA precursor was observed when expressing MPB-MpcA in the absence of any modification enzymes (**Fig. 3b**). Co-expression with MpcB and MpcC, but not with either protein alone, yielded hypermethylated core peptides, indicating that MpcBC acts as a methyltransferase pair, consistent with the genetic deletions of *mpcB* and *mpcC* above (**Fig. 3c-e**). We detected not only singly methylated species, but also peptide fragments with multiple (up to 8) methylations. The fully methylated peptide contained the same methylation pattern as clavusporin B, as judged by tandem MS analysis (**Table S6**). To probe the sequence of methylation events, the single-methylated species were analyzed by tandem HR-MS; the results pointed to a mixture of mono-methylated species, suggesting a lack of order of β-methylation by MpcBC.

We next explored the role of the MpcA leader peptide in post-translational modification. The first 17 residues of MpcA were removed from MBP-MpcA, yielding MBP-MpcA_(−1)-21_, in which the Arg at the (−1) position was retained for trypsin digestion. The truncated MBP-MpcA_(−1)-21_ was co-expressed with MpcB alone or MpcBC following the same procedure. No methylation was detectable in the absence of MpcC (**Fig. 3f**). Only a small fraction (<10%) of leader-less MpcA was mono-methylated in the presence of MpcBC, showing that the reactivity of MpcBC is largely dependent on the MpcA leader peptide (**Fig. 3g**).

The leader-peptide dependence of MpcBC suggests that methylation on MpcA precedes peptidolysis. Two separate hydrolysis events are required to remove the leader and follower peptides and thus generate the mature product. As both cleavages occur *C*-terminally to Arg, it is possible that the only peptidase encoded in the BGC, MpcP, performs these reactions. To test this idea, His_6_-tagged MpcP and NusA-His_6_-tagged MpcA were produced recombinantly in *E. coli.* Incubation of the purified proteins yielded a single product [M+3H]^3+^ with *m/z* of 758.7177, which matched MpcA_1-21_ ([M+3H]^3+^, calc. 758.7137, 5.3 ppm) (**Extended Fig. 4a**). MpcP activity is unlikely to be affected by the NusA tag, as trypsin was able to hydrolyze both upstream and downstream of the core peptide (**Extended Fig. 4b**). Moreover, when modified MBP-tagged MpcA obtained from co-expression with MpcBC was treated with MpcP, only removal of the leader peptide, but not the follower, was observed, yielding MpcA_1-21_ with 0-8 methylations (**Extended Fig. 4c**). These observations show that MpcP is a site-specific protease that removes the leader peptide from MpcA. We postulate that the acidic follower of MpcA solubilizes the peptide after extensive methylations in the core region. Once the modified MpcA_1-21_ peptide(s) are transported outside of the cell by MpcT, a yet-unknown extracellular protease removes the follower. Together, these observations allow us to propose a biosynthetic pathway of clavusporin (**Fig. 3h**).

### Mpc-type BGCs are widespread in Actinobacteria

Given the significant role the *mpc* operon in *S. clavuligerus* and *S. ghanaensis*, we explored its prevalence bioinformatically using MpcB (WP_003958123.1) as a query in protein BLAST searches (**Fig. 4a**). The first 500 non-redundant proteins were subjected to genomic neighboring analysis using RODEO^41^. MpcB-homologs situated in gene clusters that encode upstream MFS transporters and S9 peptidases as well as downstream DUF5825 proteins were selected, leading to the identification of 229 distinct *mpc* BGCs, all from Actinobacteria with identical genomic contexts. It is, therefore, reasonable to assume that these share analogous functions and generate mature products that are similar to clavusporins.

**Fig. 4.**
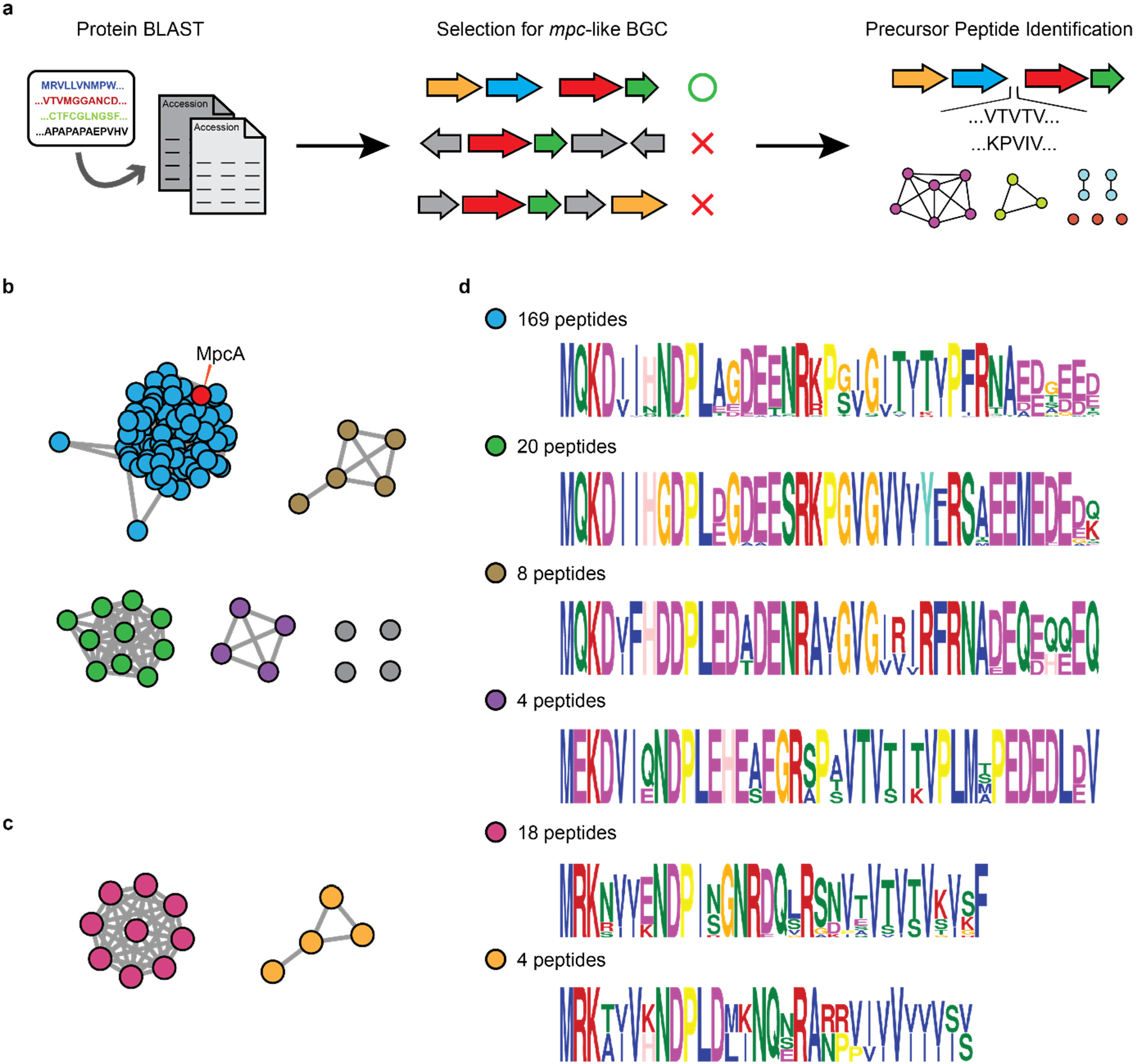
*mpc*-type BGCs are widespread in actinobacteria. **a**, Workflow for identification of *mpc* clusters and their putative precursor peptides. **b-c**, SSN of ‘long’ precursor peptides with acidic follower peptides (**b**) and ‘short’ precursor peptides without acidic follower peptides (**c**). Each node represents a unique BGC and the lines connecting them indicate sequence similarity in the precursor peptide (MpcA). The *S. clavuligerus* MpcA node is labeled in red. **d**, Logo plot for precursor peptides in each subfamily shown in **b** and **c**. The precursor peptides are color-coded to the BGCs in panels **b** and **c**, and the number of representatives for each cluster is shown.

We next identified precursor peptides for these BGCs using three criteria: Location of the ORF between MpcP and MpcB, high GC content, and a peptide sequence rich in Val, Ile, Thr and/or Pro residues. We were able to find a single precursor peptide for each of the 229 BGCs, except for one, which failed due to poor genome sequencing quality. Most of the peptides resemble MpcA, consisting of leaders, hydrophobic cores, and acidic followers. However, we also identified several (22 out of 228) short precursor peptides lacking the acidic follower sequence. Coincidently, the MpcA-like ‘long’ peptides are mostly from *Streptomyces*, with 2 exceptions out of 206 peptides, while the truncated precursors are mostly found in rare actinomycetes (**Table S7**).

To further group the identified *mpc* clusters, the 228 precursors were subjected to sequence similarity network (SSN) analysis^42^. As the algorithm is especially sensitive to sequence length, the long and short precursors were separated into two networks (**Fig. 4b-c**). This analysis organized the long precursors into four subfamilies with multiple members and four singletons. BGCs containing the short precursor were organized into two subfamilies. For each, a conserved precursor peptide motif was generated using the MEME suite (**Fig. 4d**). The majority of the long peptides (169 out of 206) cluster with MpcA, featuring two conserved Arg residues for peptidase recognition, and 2 conserved Pro residues for di-methylations. Although the number of Pro residues in the putative core regions vary among the long peptides, all have the first Arg residue for MpcP-type processing. In contrast, the second Arg in MpcA is not conserved, again pointing to a separate enzyme for follower removal. For the short precursors, the Arg residue upstream the hydrophobic core is conserved, while Pro is no longer present in the core peptides. Instead, the short peptides are rich in Val/Ile residues, with one group having an alternating pattern of Val/Ile, and another group bearing seven consecutive Val/Ile residues. These analyses annotate >200 BGCs for which the underlying chemistry, biosynthesis, enzymology and function can now be explored.

### Conclusions

It is by now near trivial to find uncharacterized BGCs in microbial genomes^16,43^. The much harder proposition is identifying the products of these clusters, and clavusporins demonstrate the challenges associated with this process as we needed to perform a genetic promoter swap, codon optimization, and unusual extraction protocols to uncover the product of the *mpc* cluster. The structure and biosynthesis of clavusporin are highly unusual. The latter not only revealed MpcBC as a novel iterative β-methyltransferase pair but also highlighted both the promiscuity and selectivity of the reaction. On one hand, MpcBC can recognize four different residues, but on the other, it consistently methylates at the β-carbon. MpcC may serve as a scaffolding domain, perhaps a RiPP Recognition Element (RRE) for B12-rSAM enzymes^44^. The reconstitution of MpcBC catalysis in *E. coli* now allows further functional studies with this heterodimer.

To the best of our knowledge, clavusporins join only polytheonamide as heavily *C*-methylated peptides^31,35^. A common approach used by nature to reduce the polarity of peptides is attachment of fatty acyl groups at the peptide *N*-terminus^45^. Installation of a bevy of individual methyl groups on amino acid side chains, as seen in clavusporins and polytheonamide, is an alternative strategy to elevate hydrophobicity. Structure and function go hand-in-hand, and the amphiphilic structure of clavusporin is well-suited for its function in lowering the surface tension of water and allowing bacteria to raise aerial structures. It has long been known that removal of the large plasmid in *S. clavuligerus* leads to a bald phenotype. Our results provide a mechanistic rationale for this observation. Clavusporins, if immobilized with a certain conformation, could be more effective as surfactants compared to free molecules in the droplet tests. The exact mechanism of deployment of clavusporins, whether they function solely as free surfactants or in addition have specific molecular targets on the cell surface, and whether they assemble on the cell wall and interact with chaplins and rodlins remain to be determined. Sporulation is a complex and coordinated process involving a number of regulators, notably the autoinducer ɣ-butyrolactone^4^. In this regard, how clavusporin connects with other regulatory pathways and known processes in sporulation provides a rich area of future research. It is also highly likely that yet-unknown natural products act as surfactants in sporulation, including those in the family of >200 *mpc*-like BGCs that we annotate, and these offer intriguing avenues for further study at the intersection of microbial development, natural product chemistry, and enzymology.

## Extended Figures

**Extended Fig. 1.**
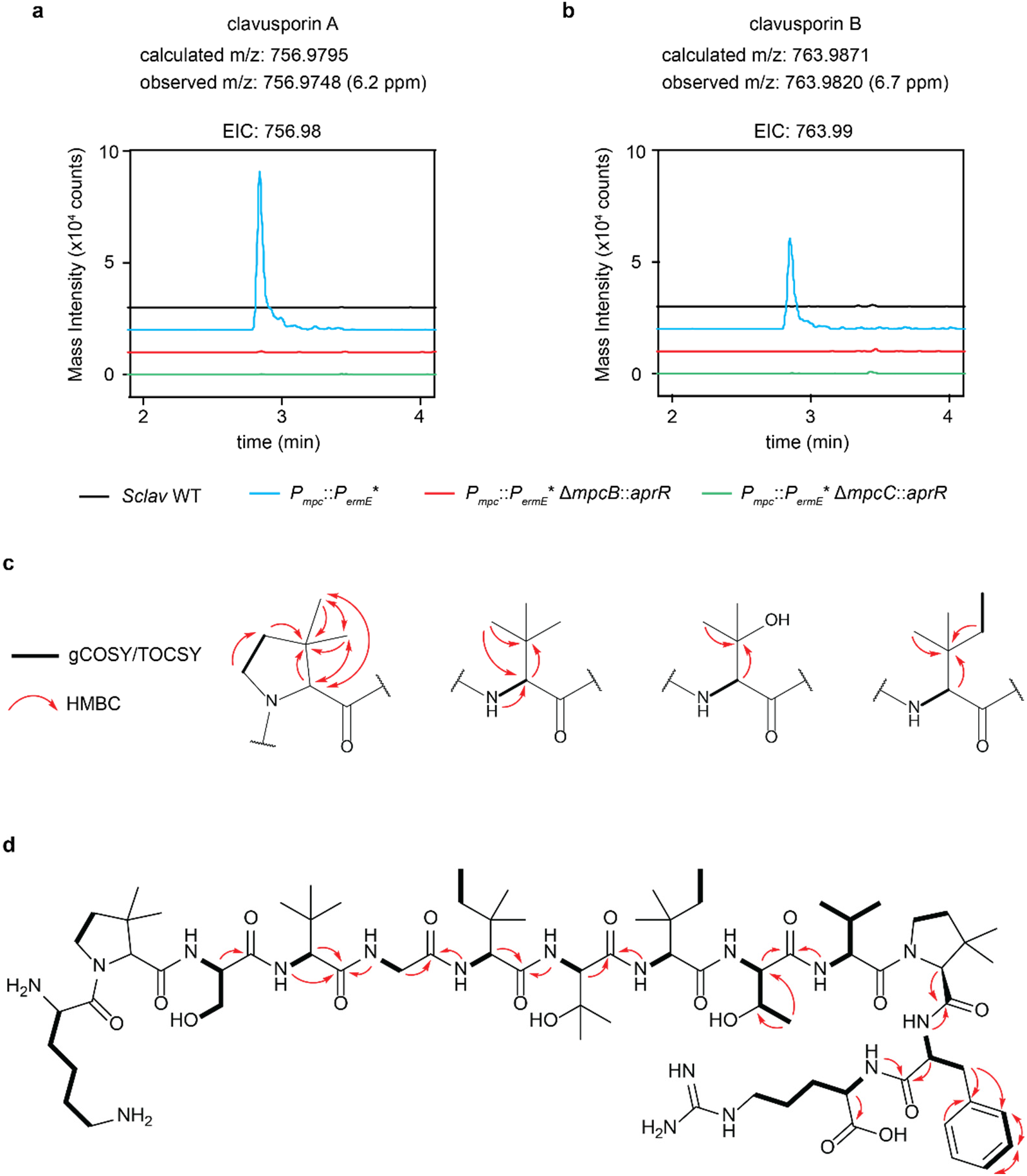
Structural elucidation of clavusporins. **a-b**, UPLC-coupled HR-MS analysis of extracts of WT *S. clavuligerus* and the deletion mutants indicated. Shown are extracted ion chromatograms of clavusporin A (**a**) and B (**b**). The traces are offset on the y-axis for clarity. **c**, Key NMR correlations used for determination of post-translational methylations in clavusporin B. **d**, Key NMR correlations used for determination of the overall structure of clavusporin B.

**Extended Fig. 2.**
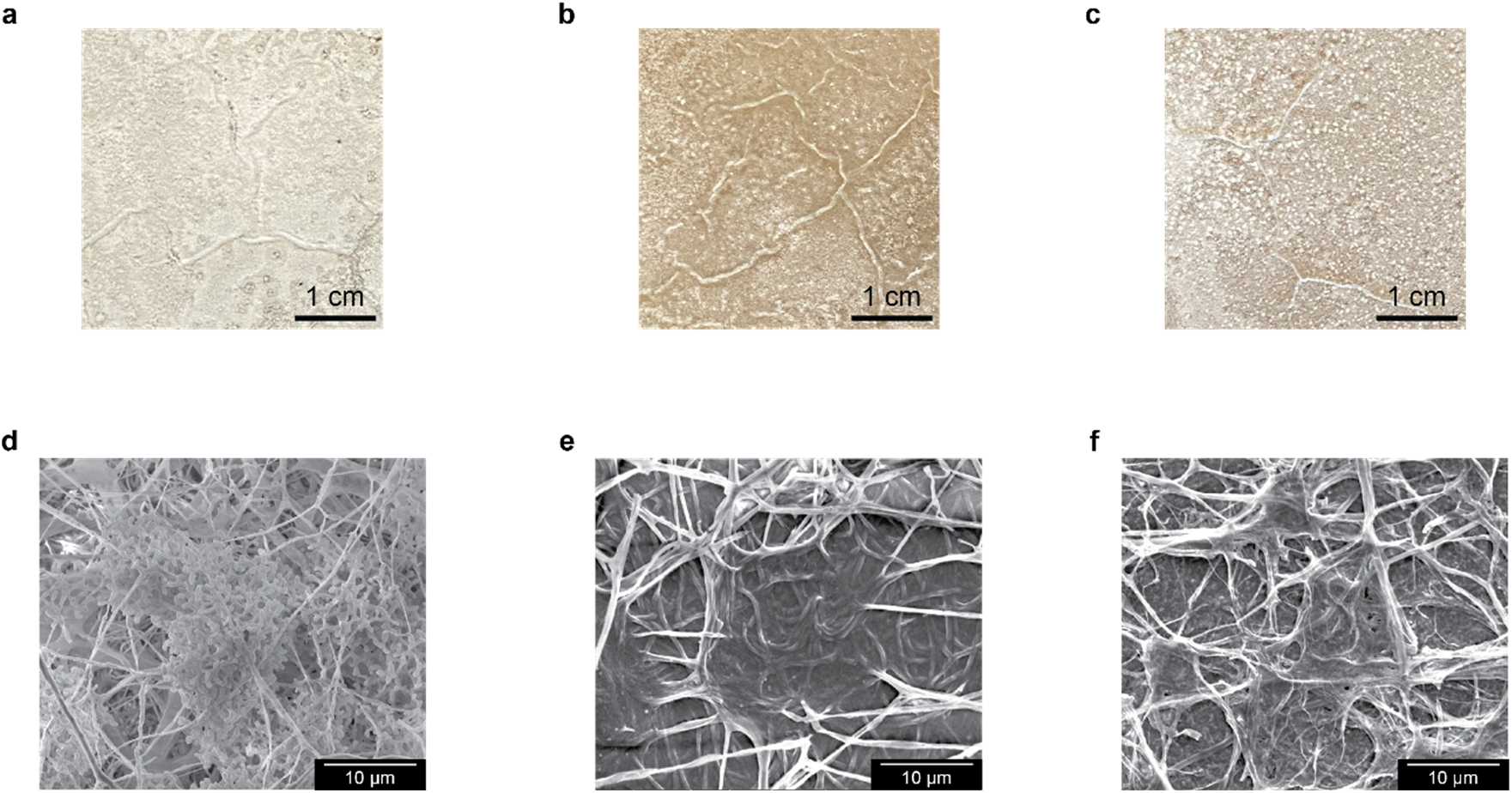
Phenotypic differences of WT *S. clavuligerus* and *mpc* mutants. **a-c**, Images of WT *S. clavuligerus* (**a**), Δ*mpcB* (**b**) and Δ*mpcC* (**c**) growing on GYM agar for 10 days. Note the green spores produced by the WT. **d-f**, SEM images of WT (**d**), Δ*mpcB* (**e**) and Δ*mpcC* (**f**), which confirm sporulation in WT, and minimal aerial hyphae growth in mutants. Each experiment was repeated in three biological replicates, and representative results are shown.

**Extended Fig. 3.**
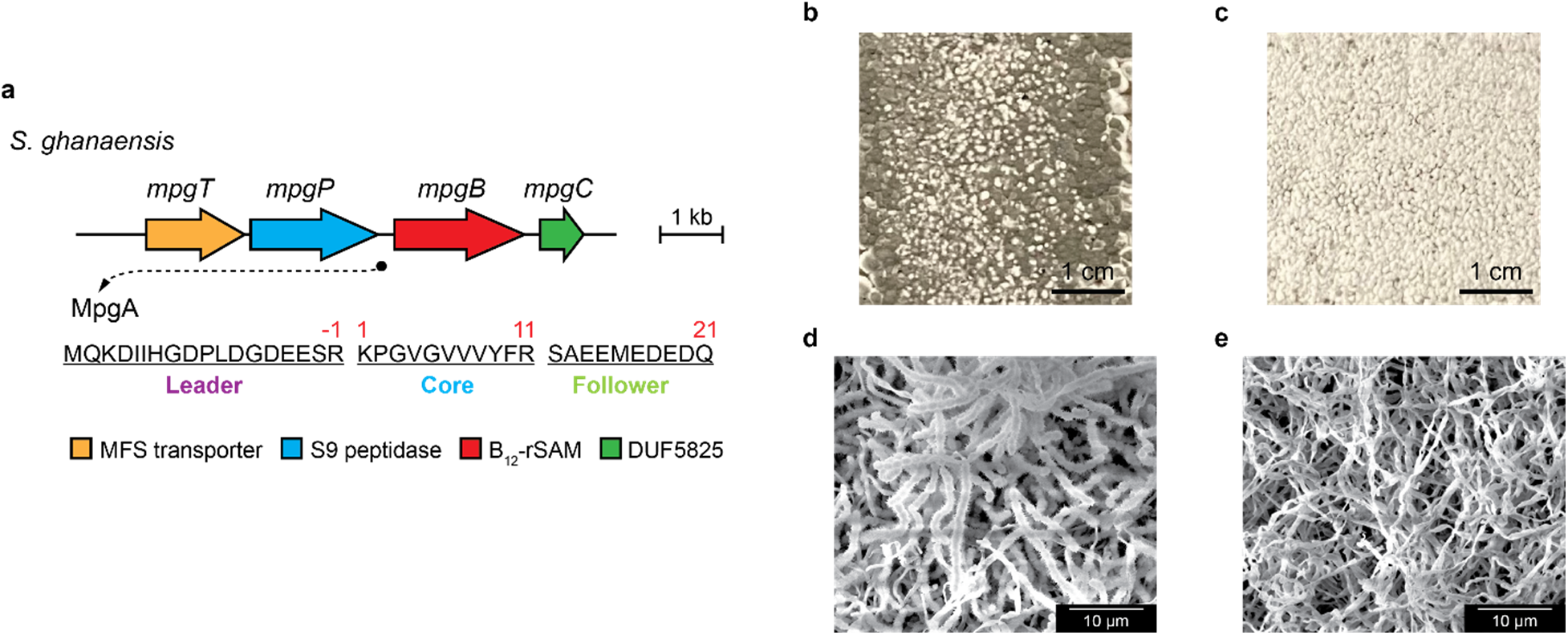
Phenotypic differences between WT *S. ghanaensis* and the *mpgB* mutant. **a**, The orthologous *mpg* gene cluster in *S. ghanaensis*. The gene locus of *mpgA* is marked by a black circle, and its protein sequence is shown. **b-c**, Images of WT *S. ghanaensis* (**b**) and Δ*mpgB S. ghanaensis* (**c**) growing on SFM agar for 3 days. Note the robust production of dark grey spores by the WT. **d-e**, SEM images of WT (**d**) and Δ*mpgB S. ghanaensis* (**e**), which show typical formation of *Streptomyces* spore chains in WT. Each experiment was repeated in three biological replicates, and representative results are shown.

**Extended Fig. 4.**
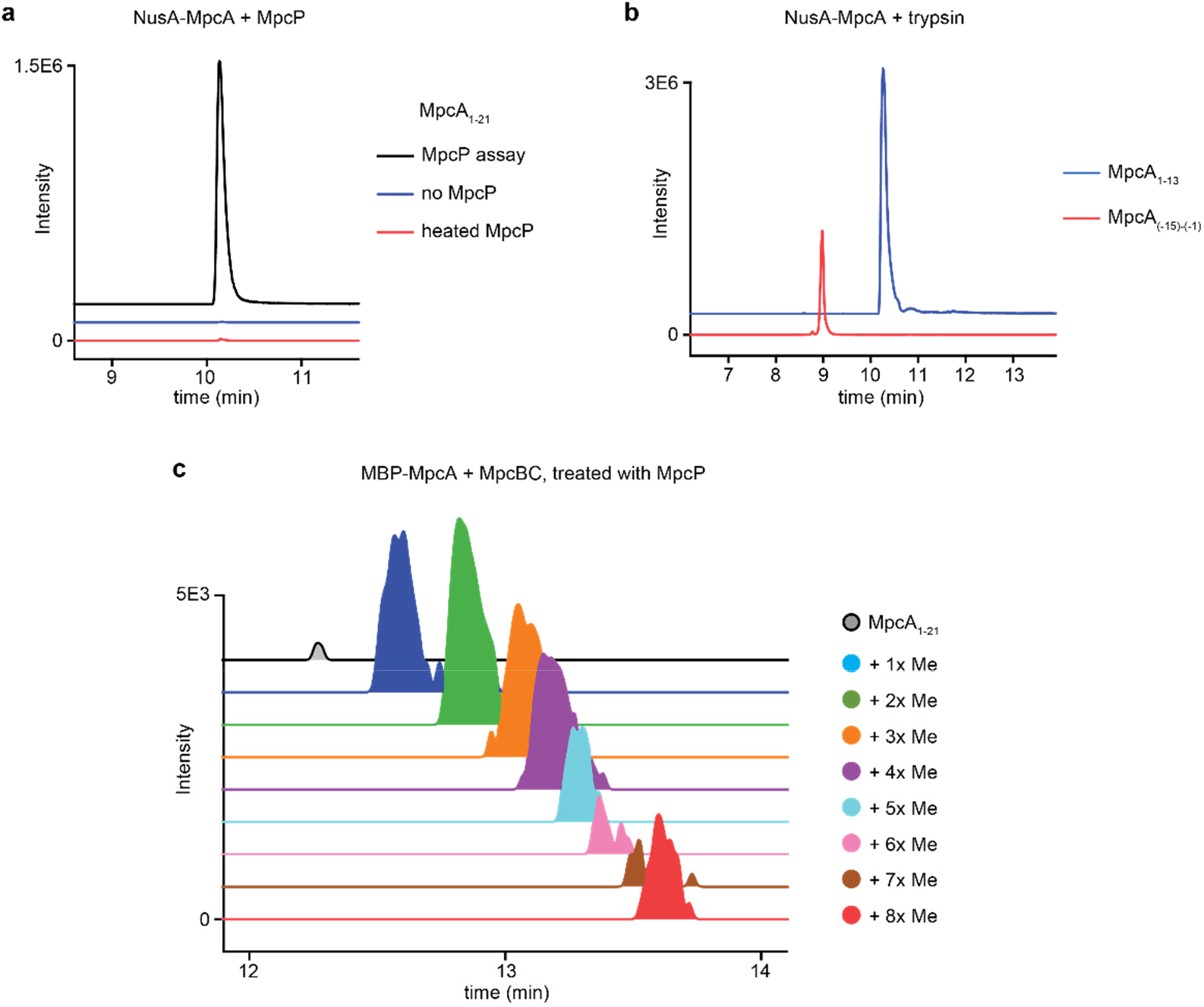
MpcP is a site-specific leader protease. **a**, UPLC-HR-MS analysis of the MpcP reaction with NusA-MpcA. Heat-inactivated MpcP and a no-enzyme reaction were used as controls. Shown are extracted ion chromatograms of MpcA_1-21_. **b**, UPLC-HR-MS analysis of trypsin-digested NusA-MpcA as a positive control of MpcP assay. Shown are extracted ion chromatograms of two tryptic fragments of MpcA, MpcA_1-13_ (blue) and MpcA_(−15)-(−1)_ (red). **c**, UPLC-HR-MS analysis of MpcP-treated MBP-MpcA fusion, which was co-expressed with MpcBC in *E. coli*. Shown are extracted ion chromatograms of MpcA_1-21_ with different methylation states. The traces are offset on the y-axis for clarity.

## Methods

### Bacteria strains and growth

*Streptomyces clavuligerus* ATCC 27064 was obtained from ATCC. *S. clavuligerus s*trains were cultured in TSB liquid medium [3% (w/v) tryptone soy broth] for general purposes. *S. clavuligerus* cultures were plated on GYM agar for sporulation [0.4% (w/v) D-glucose, 0.4% (w/v) yeast extract, 1% (w/v) malt extract, 0.2% (w/v) calcium carbonate, 1.2% (w/v) agarose, pH=7.2]. For production of clavusporin, the engineered *S. clavuligerus* strain was cultured in MYM medium [0.4% (w/v) D-maltose, 0.4% (w/v) yeast extract, 1% (w/v) malt extract]. *E. coli* strains were cultured in LB medium or LB agar with the appropriate antibiotics. Antibiotics used were ampicillin (Amp, 100 μg/mL), apramycin (Apr, 50 μg/mL), kanamycin (Kan, 50 μg/mL), chloramphenicol (Cm, 25 μg/mL), thiostrepton (Tsr, 20 μg/mL) and spectinomycin (Spec, 100 μg/mL).

### Genetic manipulation of *Streptomyces* by Conjugation

Plasmid pJTU1289 (Amp^R^, Tsr^R^) was used for the gene knockouts (*aprR* insertion) of *S.clavuligerus* and *S. ghanaensis.* Plasmid pKC1139 (Apr^R^) with a temperature-sensitive *ori* was used for the marker-less promoter exchange for *S. clavuligerus.* In brief, ∼2 kb fragments upstream and downstream of the targeted region were cloned, and inserted into the knockout plasmid, flanking the desired insertion sequence (e.g. antibiotic marker). Constructions of the knockout plasmids were completed in *E. coli* DH5α cells (Supplementary Material), and chemically transformed into methylation-deficient *E. coli* ET12567 (Cm^R^) carrying pUZ8002 (Kan^R^). The donor *E. coli* strain was then cultured in 25 mL of LB medium supplemented with antibiotics to an OD_600_ of 0.5-0.7. Cells were collected by centrifugation (4000g, 10 min), washed with antibiotic-free LB medium (10 mL, twice), and resuspended in LB medium for later use. *Streptomyces* spores were collected from agar plates by scratching the cell surface with cotton swabs and suspended in water. Excess mycelial cells were removed through crude cotton filtration. Spores remaining in the filtrate were then spun down by centrifugation, washed with 1 mL of TES buffer (50 mM TES, pH=8.0), and then resuspended in 500 μL of TES buffer. Spores were heat-shocked for 10 min in a 50°C water bath, subsequently incubated on ice for 5 min, and then mixed with 500 μL of 2x spore-activation medium [1% (w/v) yeast extract and 1% (w/v) casein hydrolysate] and calcium chloride (5 mM, final concentration). The spores were then incubated at 37°C, 200 rpm for germination for 2-3 hours, then collected by centrifugation, and resuspended in LB medium. Spore suspensions were mixed thoroughly with *E. coli* (donor/acceptor 1:1 to 2:1), plated on selected agar plates, and incubated at 30°C for 16-18 hours. Exconjugants were selected by plating Apr and trimethoprim (50 μg/mL) onto plates, and colonies emerged on plates after incubation at 30°C for 2-4 days. For gene knockouts, double-crossover colonies that were sensitive to Tsr, but resistant to Apr were selected and verified by PCR. For generating the *S. clavuligerus* engineered strain, exconjugants were first cultured in TSB with Apr at 30°C, then without Apr in TSB at 37°C for plasmid integration, and then selected with Apr again at 30°C to obtain single-crossover cells. The single-crossover cells were then plated on antibiotic-free GYM plates for single colony selection. Colonies sensitive to Apr were picked and tested with PCR for successful promoter replacement.

### Scanning Electron Microscopy

*Streptomyces* cells grown on agar plates were cut into cubes, sliced into a thin layer, and then transferred to aluminum SEM stubs (Electron Microscopy Sciences, #75220) covered with double-sided carbon tape. Samples were coated with a 3-5 nm layer of gold on a VCR IBS/TM 200S ion beam sputterer and imaged on a Quanta 200 FEG environmental-SEM in high vacuum mode.

### Detection of clavusporins from *S. clavuligerus P_mpc_*::*P_ermE_*\*

*S. clavuligerus* strains were cultured in TSB medium for 2 days at 30°C/200 rpm, and then inoculated in a 125 mL Erlenmeyer flask containing 25 mL of MYM medium (0.02% w/v), supplemented with CoCl_2_ꞏ6H_2_O (2 μg/mL, final concentration). Cells were then cultured at 30°C, 250 rpm for 6 days. After growth, cells were pelleted by centrifugation at 4000g for 20 min, resuspended in 15 mL of 1% (w/v) SDS solution, and heated for 1 hour in a 55°C water bath. The supernatants from SDS extraction were analyzed by MALDI-TOF mass spectrometry. Samples were mixed with matrix [α-cyano-4-hydroxycinnamic acid in 50% MeCN in H_2_O+0.1%TFA, 10 mg/mL] in 1:1 ratio and spotted on a ground steel 384-well target plate. MALDI-MS analysis was performed on a Bruker UltrafleXtreme TOF/TOF mass spectrometry in reflector/positive mode. For UPLC-coupled high-resolution (HR)-MS analysis, supernatants from SDS extractions were acidified with 1%(v/v) TFA and then extracted with ethyl acetate (1:1) twice. The organic phase was then combined, dried *in vacuo*, resuspended in MeOH, and subjected to UPLC-MS analysis performed on an Agilent 6546 Accurate Mass Quadrupole Time-of-Flight (Q-TOF) with a 1290 Infinity II Series LC system. LC separation was carried out on an Eclipse Plus C18 column (Agilent, 1.8 μm, 2.1 x 50 mm), with H_2_O and MeCN as mobile phases (+0.1% formic acid) at a flow rate of 0.5 mL/min. The LC method started isocratically at 10% MeCN for 0.5 min, followed by a gradient from 10% to 100% MeCN over 4.5 min, then finished with 3 min of isocratic 100% MeCN.

### Isolation of clavusporins

*S. clavuligerus P_mpc_*::*P_ermE_*\* spore stock was inoculated into a 14-mL culture tube containing 3 mL TSB medium, and cultured at 30°C, 250 rpm for 2 days. The culture was then diluted 1:100 into two 125 mL Erlenmeyer flasks containing 25 mL of TSB medium, and cultured at 30°C, 250 rpm for 2 days. The resulting cultures were used for large-scale growth. Cells were collected by centrifugation at 4000g, resuspended in MYM medium, and inoculated into 8 x 2.8-L baffled Fernbach flasks, each containing 1 L of MYM medium (0.02% w/v), supplemented with CoCl_2_ꞏ6H_2_O (2 μg/mL, final concentration). After growth at 30°C, 200 rpm for 6 days, cells were spun down at 9000 rpm, and transferred to 8 x 1 L Erlenmeyer flasks containing 600 mL of 1% SDS solution. Cell suspensions in SDS solution were warmed for 1 hour in a 55°C water bath with occasional shaking, and then stored at 4°C overnight for the precipitation of insoluble proteins. The cooled cell suspensions were then centrifuged at 9000 rpm for 90 min. The subsequent cell-free supernatants were acidified with 1% (v/v) TFA and extracted with EtOAc (1:1) twice. Liquids from the organic phase were combined, treated with sodium sulfate, and concentrated by rotary evaporation. The concentrated EtOAc extract was an oily dark brown substance; it was lyophilized to remove residual water and TFA. The resulting material was resuspended with a solution of dichloromethane (DCM)/MeOH (50:1) and loaded onto a normal phase silica column (Fisher Scientific, 230-400 mesh, grade 60). The column was eluted with DCM/MeOH 50:1, DCM/MeOH 20:1, DCM/MeOH 10:1, DCM/MeOH 5:1, DCM/MeOH 1:1, MeOH, MeOH/H_2_O 4:1 and MeOH/H_2_O 2:1. Clavusporins were detected in the last three fractions, which were then combined, and dried *in vacuo*. The material was then resuspended in 70% MeCN in H_2_O (+0.1% TFA), and further purified on an Agilent analytical HPLC system, equipped with a 1260 Infinity pump, an automatic liquid sampler, a temperature-controlled column compartment, a photodiode array detector, and an automated fraction collector. The material was repeatedly injected into an analytical Eclipse XDB-C8 column (Agilent, 5 μm, 4.6 x 150 mm), operating at a flow rate of 0.5 mL/min with a mobile phase of H_2_O and MeCN (+0.1% TFA). Elution was carried out isocratically at 25% MeCN for 5 min, followed by a gradient of 25-50% MeCN over 25 min, a gradient of 50-100% MeCN in 3 min and lastly an isocratic wash of 100% MeCN for 12 min. Fractions containing clavusporins (20-26 min), as detected by MALDI-TOF MS, were pooled, dried, and subjected to NMR analysis.

### Surface tension measurement

Compounds were dissolved in 10% DMSO (in water) for surface tension analysis on a Rame-Hart contact angle goniometer (Model 90-U3-Pro). Photos of hanging droplets (7-10 μL) were taken and analyzed by the DROPimagePro software using Surface Tension Tool. For each concentration, three technical replicates were performed.

### Chemical complementation of clavusporins

*S. clavuligerus ΔmpcB* and *ΔmpcC* knockout strains were cultured in TSB medium at 30°C, 250 rpm for 2 days, and then plated onto GYM agar plates to form an even cell lawn. Cells were first incubated at 30°C for 2 days to enable growth of substrate mycelium. Then, 2 μL of compound solutions in DMSO at various concentrations were applied to the surface of growing cells. Aerial hyphae emerged after 1-2 days of chemical complementation.

### Reconstitution of the MpcBC reaction in *E. coli*

Plasmid pRSF-Duet (Kan^R^) was used for heterologous expression of precursor peptides and the corresponding MpcBC methyltransferase in *E. coli*. Briefly, MBP-fused precursor peptide was cloned into the first multiple cloning site (MCS), and modifying enzymes were cloned into the second MCS, which were both downstream of *T7* promoters (Supplementary Material). Expression plasmids were then co-transformed with pDB1282 (Amp^R^) and pBAD42-*btuCEDFB* (Spec^R^) into *E. coli* BL21(DE3) by electroporation. Colonies from the transformation plates were cultured in LB medium (Kan, Amp, and Spec). Overnight cells from LB cultures were then inoculated into a 125 mL Erlenmeyer flask containing 80 mL of M9 salt-ethanolamine medium^30^ (Kan, Amp and Spec) to a final OD_600_ of 0.02, and cultured in the dark at 37°C, 180 rpm. Cells were induced with 0.2% (w/v) L-arabinose (final concentration) at an OD_600_ ∼0.4 and induced with isopropyl β-D-1-thiogalactopyranoside (IPTG) at an OD_600_ of 0.6-0.7 with a final concentration of 0.4 mM. Iron(III) chloride and L-cysteine hydrochloride were supplemented with both L-arabinose and IPTG induction, with final concentrations of 25 μM and 150 μM, respectively. After addition of IPTG, the cells were cultured in the dark at 37°C, 120 rpm for 20 hours. The cells were then harvested by centrifugation at 4°C (4000g, 10 min), and resuspended in 10 mL of Lysis Buffer A, which consisted of 25 mM Tris base, 300 mM NaCl, 10 mM imidazole, 10% glycerol, pH=7.7. The cell suspension was lysed by sonication (FB505 sonicator, Fisher Scientific) on ice for 50 seconds in 5 s on/15-s off cycles at 30% power. Two mL of the cell lysate was then centrifuged at 4°C, 13,400 rpm for 10 min, and the supernatant loaded onto a pre-equilibrated mini-Ni column (Thermo Scientific, HisPur Ni-NTA Spin Column, 0.2 mL). The Ni column was then washed with 1 mL of Buffer A, 1 mL of Wash Buffer B (25 mM Tris base, 300 mM NaCl, 50 mM imidazole, 10% glycerol, pH=7.7), and then eluted with 1 mL of Elution Buffer C (25 mM Tris base, 300 mM NaCl, 300 mM imidazole, 10% glycerol, pH=7.7). Eluates from the Ni column were combined and loaded onto a desalting column (PD MiniTrap G-10, Cytiva) to buffer exchange into Storage Buffer D, which consisted of 25 mM Tris base, 150 mM NaCl, 10% glycerol, pH=7.7, following the manufacturer’s instruction. Enriched MBP-tagged peptides (100 μL) were subsequently treated with trypsin (∼0.5 μg) in the presence of CaCl_2_ (20 mM) at 37°C overnight. The digested peptide mixture was quenched at 95°C for 5 min, diluted with 100 μL of MeOH, and subjected to LC-MS analysis, which was performed on a 6540 UHD Accurate Mass Q-tof LC-MS system (Agilent), consisting of a 1260 Infinity Series HPLC, an automated liquid sampler, a diode array detector, and a JetStream ESI source, using an analytical Jupiter C18 column (Phenomenex, 5 μm, 300 Å, 4.6 x 150 mm) with a gradient of 5% MeCN in H_2_O (+0.1% formic acid) to 95% MeCN in H_2_O (+0.1% formic acid) over 15 min.

### Protein overexpression and purification

Genes *mpcA* and *mpcP* were amplified from *S. clavuligerus* genomic DNA and cloned into the BamHI/XhoI site of plasmid pIJ-NusA (Amp^R^) and the NdeI/XhoI site of pET28b (Kan^R^) for overexpression of NusA-MpcA and His_6_-MpcP, respectively (**Table S5**). The plasmids were then transformed into *E. coli* BL21 (DE3) via chemical transformation. Colonies from the transformation plates were inoculated into LB medium (with antibiotics), and then transferred to a 125 mL Erlenmeyer flask containing 25 mL of LB as seed culture. After 16-18 hours, cells from the 25 mL culture were inoculated into a 4 L Erlenmeyer flask containing 800 mL of LB medium with an initial OD_600_ of 0.02, and cultured at 37°C, 180 rpm. Cultures were induced with IPTG (0.1 mM, final concentration) at an OD_600_ of 0.4-0.6, and then cultured at 18°C, 180 rpm for 18 hours. The cells were pelleted by centrifugation at 4°C (4000g, 20 min) and resuspended in ∼60 mL of Buffer A. These were lysed by sonication on ice for 5 min in 5 s on/15 s off cycles at 35% power. The cell lysate was then centrifuged at 4°C, 12,000 rpm for 60 min, and the supernatant loaded onto a Ni-affinity column (5 mL), which was pre-equilibrated with 50 mL of Buffer A. The column was washed with 50 mL of Lysis buffer A and subsequently 50 mL of Wash Buffer B. Overexpressed protein was then eluted with Elution Buffer C, and fractions containing the target proteins, as analyzed by SDS-PAGE gel, were combined and buffer-exchanged into Storage Buffer D using ultra-filters (Amicon). Concentrated protein solutions were then aliquoted and stored in −80°C.

### MpcP assay

Typical MpcP reactions were performed in 100 μL of Storage Buffer D with ∼20 μM of purified MpcP and ∼100 μM of NusA-MpcA. Control assays were conducted by adding heat-inactivated (95°C, 10 min) MpcP or no addition of MpcP. Assay mixtures were incubated at 30°C overnight, quenched with 2 volumes of MeOH, and centrifuged at 21,100g for 10 min. The supernatant was then subjected to HPLC-MS analysis as described above.

### Purification of the unmodified core peptide

NusA-tagged MpcA and MpcP were overexpressed and purified as described above, and a large-scale reaction (10 mL) was performed with 200 μM of NusA-MpcA and ∼50 μM of MpcP at 30°C overnight. The reaction was quenched with 2 volumes of acetonitrile, centrifuged (4°C, 4000g, 20 min), and the supernatant then transferred to a glass vial for rotary evaporation. After the solvent was removed, 100 μg of trypsin and calcium chloride (20 mM final concentration) were added to the aqueous solution for further digestion at 37°C overnight. Then 5 g of HP20 resins (Thermo Fisher) was added to the reaction mixture to absorb the desired product. The HP20 resin was then isolated, washed with 5 volumes of each water, 5% MeCN+0.1%TFA, 25% MeCN+0.1%TFA, 50% MeCN+0.1%TFA, 75% MeCN+0.1%TFA, and 100% MeCN+0.1%TFA. Fractions containing unmodified MpcA_1-13_, as judged by HPLC-MS, were pooled, dried *in vacuo*, and resuspended in 70% MeCN+0.1%TFA for further purification. The material was then purified on an Agilent analytical HPLC as described above, with an analytical Eclipse XDB-C8 column (Agilent, 5μm, 4.6x150 mm), operating at a flow rate of 0.5 mL/min with mobile phase of H_2_O and MeCN (+0.1% TFA). The elution was carried out first isocratically at 15% MeCN for 5 min, followed by a gradient of 15-40% MeCN over 25 min, a gradient of 40-100% MeCN over 3 min and lastly isoacratic elution with 100% MeCN for 12 min. Fractions containing unmodified MpcA_1-13_ (21-24 min), as judged by HPLC-MS, were pooled, dried, and subjected to NMR analysis.

### Bioinformatics

Protein BLAST was performed with the MpcB sequence and default blastp parameters on NCBI. The accession numbers of the first 500 aligned sequences were downloaded and subjected to genome neighborhood analysis using the RODEO webtool. Gene clusters encoding MFS transporters, S9 peptidase or alpha/beta hydrolase fold, B_12_-rSAM enzymes and hypothetical proteins (DUF5825) were selected manually and further analyzed for potential precursor peptides. The GC% of codon usage for different open-reading frames were analyzed by online in FramePlot. In total, 228 *mpc*-like gene clusters were identified with putative precursors, encompassing 104 unique peptide sequences. Precursor peptides are grouped into ‘long’ peptides (with acidic follower, length ≥ 37 aa) and ‘short’ peptides (without acidic follower, length ≤ 33 aa). Two groups of precursor peptides were separately input into EFI-EST for SSN analysis using option C. An alignment score threshold of 15 was chosen for ‘long’ peptides, and 8 for ‘short’ peptides. The SSNs were visualized in Cytoscape with prefused force directed layout. Conserved motif analysis of each cluster was performed on MEME suite using the GLAM2 algorithm.

## Acknowledgements

The authors thank the Seyedsayamdost lab members for helpful discussions, Dr. John Schreiber for assistance in SEM experiments, Dr. Denis Potapenko for guidance in surface tension measurement and Dr. István Pelczer for collecting NMR data. We thank the National Science Foundation (NSF CAREER Award 1847932 to M.R.S.) as well as the National Institutes of Health (grant GM140034 to M.R.S.) for financial support.

## Author Contributions

C.Z. and M.R.S. conceived of the project. C.Z. designed and performed experiments and analyzed data. C.Z. and M.R.S. wrote the manuscript.

## Disclosures

The authors declare no competing interests.

## Notes

### Competing Interest Statement

The authors have declared no competing interest.

## References

1. Clardy, J. & Walsh, C. Lessons from natural molecules. Nature 432, 829–837 (2004).

2. Salvador-Reyes, L. A. & Luesch, H. Biological targets and mechanisms of action of natural products from marine cyanobacteria. Nat. Prod. Rep. 32, 478–503 (2015).

3. Wang, R. & Seyedsayamdost, M. R. Hijacking exogenous signals to generate new secondary metabolites during symbiotic interactions. Nat. Rev. Chem. 1, 0021 (2017).

4. van der Meij, A., Worsley, S. F., Hutchings, M. I. & van Wezel, G. P. Chemical ecology of antibiotic production by actinomycetes. FEMS Microbiol. Rev. 41, 392–416 (2017).

5. Bassler, B. L. & Losick, R. Bacterially speaking. Cell 125, 237–246 (2006).

6. Hider, R. C. & Kong, X. Chemistry and biology of siderophores. Nat. Prod. Rep. 27, 637–657 (2010).

7. Khan, S. H., Ahmad, N., Ahmad, F. & Kumar, R. Naturally occurring organic osmolytes: from cell physiology to disease prevention. IUBMB Life 62, 891–895 (2010).

8. Ron, E. Z. & Rosenberg, E. Natural roles of biosurfactants. Environ. Microbiol. 3, 229–236 (2001).

9. Straight, P. D., Willey, J. M. & Kolter, R. Interactions between *Streptomyces coelicolor* and *Bacillus subtilis*: Role of surfactants in raising aerial structures. J. Bacteriol. 188, 4918–4925 (2006).

10. Bibb, M. J. Regulation of secondary metabolism in streptomycetes. Curr. Opin. Microbiol. 8, 208–215 (2005).

11. Flärdh, K. & Buttner, M. J. *Streptomyces* morphogenetics: dissecting differentiation in a filamentous bacterium. Nat. Rev. Microbiol. 7, 36–49 (2009).

12. Chater, K. F. Recent advances in understanding *Streptomyces*. F1000Res. 5, 2795 (2016).

13. McCormick, J. R. & Flärdh, K. Signals and regulators that govern *Streptomyces* development. FEMS Microbiol. Rev. 36, 206–231 (2012).

14. Tillotson, R. D., Wösten, H. A., Richter, M. & Willey, J. M. A surface active protein involved in aerial hyphae formation in the filamentous fungus *Schizophillum commune* restores the capacity of a bald mutant of the filamentous bacterium *Streptomyces coelicolor* to erect aerial structures. Mol. Microbiol. 30, 595–602 (1998).

15. Wösten, H. A. B. & Willey, J. M. Surface-active proteins enable microbial aerial hyphae to grow into the air. Microbiology 146 (Pt 4), 767–773 (2000).

16. Gavriilidou, A. et al. Compendium of specialized metabolite biosynthetic diversity encoded in bacterial genomes. Nat. Microbiol. 7, 726–735 (2022).

17. Kodani, S. et al. The SapB morphogen is a lantibiotic-like peptide derived from the product of the developmental gene *ramS* in *Streptomyces coelicolor*. Proc. Natl. Acad. Sci. USA 101, 11448–11453 (2004).

18. Willey, J., Santamaria, R., Guijarro, J., Geistlich, M. & Losick, R. Extracellular complementation of a developmental mutation implicates a small sporulation protein in aerial mycelium formation by *S. coelicolor*. Cell 65, 641–650 (1991).

19. RiPP Montalbán-López, M., et al. New developments in RiPP discovery, enzymology and engineering. Nat. Prod. Rep. 38, 130–239 (2021).

20. Álvarez-Álvarez, R. et al. A 1.8-Mb-reduced *Streptomyces clavuligerus* genome: relevance for secondary metabolism and differentiation. Appl. Microbiol. Biotechnol. 98, 2183–2195 (2014).

21. Söding, J., Biegert, A. & Lupas, A. N. The HHpred interactive server for protein homology detection and structure prediction. Nucleic Acids Res. 33, W244–8 (2005).

22. Broderick, J. B., Duffus, B. R., Duschene, K. S. & Shepard, E. M. Radical S-adenosylmethionine enzymes. Chem. Rev. 114, 4229–4317 (2014).

23. Frey, P. A., Hegeman, A. D. & Ruzicka, F. J. The Radical SAM Superfamily. Crit. Rev. Biochem. Mol. Biol. 43, 63–88 (2008).

24. Landgraf, B. J., McCarthy, E. L. & Booker, S. J. Radical S-Adenosylmethionine Enzymes in Human Health and Disease. Annu. Rev. Biochem. 85, 485–514 (2016).

25. Zhang, Q., van der Donk, W. A. & Liu, W. Radical-mediated enzymatic methylation: a tale of two SAMS. Acc. Chem. Res. 45, 555–564 (2012).

26. Bauerle, M. R., Schwalm, E. L. & Booker, S. J. Mechanistic diversity of radical S-adenosylmethionine (SAM)-dependent methylation. J. Biol. Chem. 290, 3995–4002 (2015).

27. Zhou, S. et al. Mechanistic insights into class B radical-S-adenosylmethionine methylases: ubiquitous tailoring enzymes in natural product biosynthesis. Curr. Opin. Chem. Biol. 35, 73– 79 (2016).

28. Huo, L. et al. Synthetic biotechnology to study and engineer ribosomal bottromycin biosynthesis. Chem. Biol. 19, 1278–1287 (2012).

29. Freeman, M. F. et al. Metagenome mining reveals polytheonamides as posttranslationally modified ribosomal peptides. Science 338, 387–390 (2012).

30. Crone, W. J. K. et al. Dissecting Bottromycin Biosynthesis Using Comparative Untargeted Metabolomics. Angew. Chem. Int. Ed Engl. 55, 9639–9643 (2016).

31. Freeman, M. F., et al. Seven enzymes create extraordinary molecular complexity in an uncultivated bacterium. Nat. Chem. 9, 387–395 (2017).

32. Noike, M. et al. A peptide ligase and the ribosome cooperate to synthesize the peptide pheganomycin. Nat. Chem. Biol. 11, 71–76 (2015).

33. Schmitt-John, T. & Engels, J. W. Promoter constructions for efficient secretion expression in *Streptomyces lividans*. Appl. Microbiol. Biotechnol. 36, 493–498 (1992).

34. Guijarro, J., Santamaria, R., Schauer, A. & Losick, R. Promoter determining the timing and spatial localization of transcription of a cloned *Streptomyces coelicolor* gene encoding a spore-associated polypeptide. J. Bacteriol. 170, 1895–1901 (1988).

35. Hamada, T. et al. Solution structure of polytheonamide B, a highly cytotoxic nonribosomal polypeptide from marine sponge. J. Am. Chem. Soc. 132, 12941–12945 (2010).

36. Woolfrey, S. G., Banzon, G. M. & Groves, M. J. The effect of sodium chloride on the dynamic surface tension of sodium dodecyl sulfate solutions. J. Colloid Interface Sci. 112, 583–587 (1986).

37. Higgens, C. E. & Kastner, R. E. *Streptomyces clavuligerus* sp. nov., a -Lactam Antibiotic Producer. Int. J. Syst. Bacteriol. 21, 326–331 (1971).

38. Sinner, E. K., Marous, D. R. & Townsend, C. A. Evolution of Methods for the Study of Cobalamin-Dependent Radical SAM Enzymes. ACS Bio. Med. Chem. Au 2, 4–10 (2022).

39. Lanz, N. D. et al. Enhanced Solubilization of Class B Radical S-Adenosylmethionine Methylases by Improved Cobalamin Uptake in *Escherichia coli*. Biochemistry 57, 1475–1490 (2018).

40. Blaszczyk, A. J., Wang, R. X. & Booker, S. J. TsrM as a Model for Purifying and Characterizing Cobalamin-Dependent Radical S-Adenosylmethionine Methylases. Methods Enzymol. 595, 303–329 (2017).

41. Tietz, J. I. et al. A new genome-mining tool redefines the lasso peptide biosynthetic landscape. Nat. Chem. Biol. 13, 470–478 (2017).

42. Gerlt, J. A. Genomic Enzymology: Web Tools for Leveraging Protein Family Sequence-Function Space and Genome Context to Discover Novel Functions. Biochemistry 56, 4293– 4308 (2017).

43. Blin, K. et al. antiSMASH 7.0: new and improved predictions for detection, regulation, chemical structures and visualisation. Nucleic Acids Res. (2023) doi:10.1093/nar/gkad344.

44. Burkhart, B. J., Hudson, G. A., Dunbar, K. L. & Mitchell, D. A. A prevalent peptide-binding domain guides ribosomal natural product biosynthesis. Nat. Chem. Biol. 11, 564–570 (2015).

45. Zhang, S. et al. Structural diversity, biosynthesis, and biological functions of lipopeptides from *Streptomyces*. Nat. Prod. Rep. 40, 557–594 (2023).

