## Supporting Information for "Widespread Peptide Surfactants with Post-translational *C-*methylations Promote Bacterial Development"

**Replacing $P_{mpc}$ with $P_{ermE}$* in *S. clavuligerus***

To insert $P_{ermE}$* and replace the rare TTA codon, 401 bp upstream of *mpcT* and the first 33 bp of *mpcT* were targeted for genetic replacement. ~2 kb fragments upstream and downstream of the targeted region were amplified from genomic DNA by PCR using primers *ScvIns_L_for*, *ScvIns_L_rev*, and *ScvIns_R_for*, *ScvIns_R_rev_E*, respectively (Table S1). The PCR products were ligated into EcoRV-digested pBluescript SK(-) through blunt end ligation and verified by Sanger sequencing. The right arm (downstream 2-kb fragment) was then amplified with primers *ScvIns_R_rev_X* and *ScvIns_R_RBS_MFS_for* to incorporate ribosomal binding site (RBS) and synonymous codons for the first 11 amino acids of MpcT and cloned into the NdeI/XbaI site of plasmid pIB139, which contains the sequence of $P_{ermE}$*. The right arm along with 5'-fused insertion sequence was then amplified by primers *ScvIns_R_PermE_for* and *ScvIns_R_rev_E*, ligated into pBluescript SK(-), and verified by Sanger sequencing. Three-piece DNA ligation was performed with HindIII/XbaI-excised left arm, XbaI/EcoRI-excised right arm with 5'-fused insertion, and HindIII/EcoRI-linearized plasmid pKC1139, successful plasmid construction was verified by enzymatic digestion. The engineering plasmid was then introduced into *S. clavuligerus* by conjugation (*Methods*). Double cross-over colonies were screened for susceptibility of Apr, and further tested by PCR (Figure S2).

**Generating Insertional mutants in *S. clavuligerus* and *S. ghanaensis***

The genomic DNAs of the *Streptomyces* strains were extracted using the Wizard Genomic DNA Purification Kit (Promega). ~2 kb fragments upstream and downstream of targeted genes were amplified by PCR using Q5 polymerase and primers listed in Table S5-6. The PCR products were cloned into EcoRV-digested pBluescript SK(-) through blunt end ligation, sequenced and then excised by listed restriction enzymes for further plasmid construction. The insertion fragment 773 cassette (Apr resistance gene *aac(3)-IV* fused with *oriT*) was obtained through digestion of plasmid pIJ773 by EcoRI and HindIII. All digested fragments (homolog arms, *aprR* insertion and linearized plasmid pJTU1289) were ligated by T4 DNA ligase at room temperature for 2 hours, and chemically transformed into *E. coli* DH5α cells, with addition of Amp and Apr for selection. Successful knockout plasmid constructions were confirmed by enzymatic digestion. Plasmids were introduced into streptomyces cells by conjugation (*Methods*). Double cross-over colonies were selected by Tsr susceptibility and Apr resistance and confirmed by PCR (Figures S6 and S8).

**Constructing plasmids and strains for co-expression of MBP-MpcA and modifying enzymes**

MpcA was inserted into the first multiple cloning site (MCS) of pRSF-Duet for the expression of $His_6$-MBP-tagged constructs. *MpcA* was cloned from the genomic DNA of *S. clavuligerus* using primers MpcA_for and MpcA_rev to yield plasmid plasmid *pRSF-Duet-H6MBP-MpcA* (Table S5). For co-expression of MBP-MpcA and MpcB, *mpcB* was amplified from genomic DNA and inserted into the second MCS of *pRSF-Duet-H6MBP-MpcA* between NdeI/KpnI sites, yielding plasmid *pRSF-Duet-H6MBP-MpcA+MpcB*. To co-express MpcC with MBP-MpcA and MpcB, *mpcC* was first amplified from genomic DNA using primers *MpcC_for* and *MpcC_rev*, and then amplified using primers *MpcC_for_GA* and *MpcC_rev_GA* for incorporation into XhoI site of *pRSF-Duet-H6MBP-MpcA+MpcB* through Gibson assembly. For co-expression of MBP-MpcA and MpcC, *mpcC* was amplified using primers *MpcC_for_NdeI* and *MpcC_rev* and inserted into the second MCS of of *pRSF-Duet-H6MBP-MpcA* between NdeI/XhoI sites. The co-expression plasmids were then co-transformed with plasmids pDB1282 ($Amp^R$) and pBAD43-BtuCEDFB ($Spec^R$) into *E. coli* BL21 (DE3) cells by electroporation.

For electroporation, overnight *E. coli* BL21 (DE3) culture was inoculated into a 250-mL Erlenmeyer flask containing 50 mL of LB medium with an initial $OD_{600}$ below 0.05, and grew at 37°C, 200 rpm until $OD_{600}$ reached around 0.6. After growth, cells were collected by centrifugation at 4°C, 4,000g for 5 min, washed with 10 mL of cooled water twice at 4°C, and washed with 10 mL of cooled 10% glycerol (in water) twice at 4°C. Cells were then resuspended in 1 mL of 10% glycerol, aliquoted into 1.5-mL Eppendorf tubes (on ice) in 100-µL volumes. For each transformation, approximately 100 ng of each plasmid was added to 100-µL cell suspension, and mixed gently. Cells were incubated with plasmids on ice for 15 min, transferred to a pre-chilled electroporation cuvette (0.1 cm, Bio-Rad), and electric shocked on a Gene Pulser Xcell Electroporation Systems (Bio-Rad) using a pre-set *E. coli* protocol (exponential decay, voltage=1800 V, capacitance=25 µF, resistance=200 Ω). After electroporation, cuvettes were placed on ice, and cells were suspended with 800 µL of LB medium, transferred to Eppendorf tubes, incubated at 37°C, 200 rpm for 30 min, and plated on LB agar plates containing Amp, Kan and Spec. After incubation at 37°C overnight, colonies on the plates were picked, and cultured for co-expression experiments.

**Table S1.** Primers used for genetic manipulation of *S. clavuligerus*. Sequences are shown in the 5'-3' direction. Recognition sites of restriction enzymes are underlined.

| Primer | Enzyme | Sequence |
|---|---|---|
| Scv_ΔmpcB_L_for | SpeI | ATATATACTAGTACTGCTGGACGTGCCTTCCGGG |
| Scv_ΔmpcB_L_rev | EcoRI | ATATATGAATTCATTGGCGTGCAGGACTTCGGCCG |
| Scv_ΔmpcB_R_for | HindIII | ATATATAAGCTTTGTGCAGCGGCTCCTCGACTCC |
| Scv_ΔmpcB_R_for | XhoI | ATATATCTCGAGAGTACAAGGAGCTGGGCACGGAGC |
| Scv_ΔmpcC_L_for | SpeI | ATATATACTAGTTGACGCTCGCGCTGCACAAGACG |
| Scv_ΔmpcC_L_rev | EcoRI | ATATATGAATTCAGGCGCAGCTCCCACTCGACG |
| Scv_ΔmpcC_R_for | HindIII | ATATATAAGCTTTTCACCGCCGACGAGCCGCTG |
| Scv_ΔmpcC_R_for | XhoI | ATATATCTCGAGAGATGGCCGTCGACTGGGTCGTG |
| ScvIns_L_for | HindIII | ATATATAAGCTTTGACCTGTGCATTGAGCCTGCACTCCG |
| ScvIns_L_rev | XbaI | ATATATTCTAGATGGGAACGGTGGTGGTGAGGGTACG |
| ScvIns_R_for | EcoRI | ATATATGAATTCTCGTTTCGTCCCCACCACACTTTCTGC |
| ScvIns_R_rev_E | EcoRI | ATATATGAATTCTGATGCCGGTGAGCGTGGCATGG |
| ScvIns_R_rev_X | XbaI | ATATATTCTAGATGATGCCGGTGAGCGTGGCATGG |
| ScvIns_R_RBS_MFS_for | NdeI | ATCATATGaacAGGAGGccccagATGACCTCGCTGCAGG<br>CGACGCTGGGGAGCAGGtcgtttcgtccccaccacactttctgc |
| ScvIns_R_PermE_for | XbaI | ATATATTCTAGAcatgcgagtgtccgttcgagtggcg |
| ScvIns_check_for | - | TTCGCCCCGTCGTACCGCTCC |
| ScvIns_check_rev | - | AACGGTCGTGCGGGTGACTCTGACC |

**Table S2.** NMR assignments for clavusporin B in DMSO-$d_6$.

| Residue | | δH[a] | δC[b] | Residue | | δH[a] | δC[b] |
|---|---|---|---|---|---|---|---|
| Lys (1) | CO | - | 169.5 | mThr (7) | CO | - | 173.2 |
| | N | n.d. | - | | N | 7.78 | - |
| | α | 4.44 | 59.9 | | α | 4.20 | 59.1 |
| | β | 1.83, 2.03 | 29.5 | | β | - | 66.8 |
| | γ | 1.26, 1.77 | 22.5 | | γ | 1.05 | 29.7 |
| | δ | 2.87, 3.10 | 38.7 | Thr (9) | CO | - | 172.2 |
| | ε | 3.14, 3.60 | 47.3 | | N | 7.73 | - |
| dmPro (2/11) | CO | - | 171.3 | | α | 4.08 | 59.0 |
| | N | - | - | | β | 4.07 | 66.4 |
| | α | 4.04 | 58.9 | | γ | 1.03 | 20.5 |
| | β | - | 30.72 | Val (10) | CO | - | n.d. |
| | β-$CH_3$ | 0.78, 0.84 | 18.5, 19.6 | | N | 8.46 | - |
| | γ | 1.39 | 33.0 | | α | 4.07 | 51.3 |
| | δ | 3.36 | 45.4 | | β | 1.53, 1.66 | 29.5 |
| Ser (3) | CO | - | 171.0 | | γ | 0.83, 0.86 | 22.1 |
| | N | 8.22 | - | Phe (12) | CO | - | 172.6 |
| | α | 4.34 | 55.7 | | N | 8.13 | - |
| | β | 3.62 | 61.7 | | α | 4.47 | 54.1 |
| | Oγ | 4.98 | - | | β | 2.91, 3.02 | 36.5 |
| mVal (4) | CO | - | 167.3 | | γ | - | 137.8 |
| | N | 8.07 | - | | δ | 7.28 | 129.8 |
| | α | 4.18 | 55.8 | | ε | 7.27 | 128.6 |
| | β | - | 39.5 | | ζ | 7.17 | 130.7 |
| | γ | 0.86 | 24.6 | Arg (13) | CO | - | 172.1 |
| Gly (5) | CO | - | 171.9 | | N | 8.02 | - |
| | N | 8.14 | - | | α | 4.21 | 53.2 |
| | α | 3.71 | 55.7 | | β | 2.14, 2.32 | 30.4 |
| mIle (6/8) | CO | - | 173.1 | | γ | 1.81, 1.93 | 27.7 |
| | N | 8.17/8.04 | - | | δ | 3.37 | 41.4 |
| | α | 4.18/4.17 | 52.2 | | | | |
| | β | - | 39.8 | | | | |
| | β-$CH_3$ | 0.88/0.80 | 23.5 | | | | |
| | γ | 1.59 | 24.0 | | | | |
| | δ | 0.60 | 9.5 | | | | |

[a]800 MHz; [b]Determined by HSQC and HMBC at 800 MHz; n.d., not detected.

**Table S3.** NMR assignments for unmodified $MpcA_{1-13}$ in DMSO-$d_6$.

| Residue | | δH[a] | δC[b] | Residue | | δH[a] | δC[b] |
|---|---|---|---|---|---|---|---|
| Lys (1) | CO | - | n.d. | Ile (6/8) | CO | - | 171.0/169.6 |
| | N | 8.15 | - | | N | 7.70/7.85 | - |
| | α | 4.13 | 50.4 | | α | 4.30/4.34 | 59.1/58.3 |
| | β | 1.71 | 29.2 | | β | 1.72/1.68 | 36.6/36.8 |
| | γ | 1.44 | 20.4 | | β-$CH_3$ | 0.80/0.82 | 15.0 |
| | δ | 1.54 | 26.4 | | γ | 1.07/1.04 | 24.0 |
| | ε | 2.75 | 28.3 | | δ | 0.78/0.79 | 10.8 |
| | Nζ | 7.77 | - | Thr (7/9) | CO | - | n.d. |
| Pro (2/11) | CO | - | 171.3 | | N | 7.90/7.99 | - |
| | N | - | - | | α | 4.25/4.22 | 57.7/58.0 |
| | α | 4.49 | 59.2 | | β | 3.94/3.91 | 66.4/66.3 |
| | β | 1.85, 2.12 | 29.0 | | γ | 0.98/1.00 | 19.6 |
| | γ | 1.84, 1.92 | 24.3 | Phe (12) | CO | - | n.d. |
| | δ | 3.46, 3.67 | 46.8 | | N | 7.84 | - |
| Ser (3) | CO | - | 169.8 | | α | 4.48 | 53.5 |
| | N | 8.20 | - | | β | 2.85, 3.02 | 36.9 |
| | α | | 54.9 | | γ | - | 137.7 |
| | β | 3.56, 3.64 | 61.3 | | δ | 7.26 | 129.2 |
| | Oγ | n.d. | - | | ε | 7.24 | 127.9 |
| Val (4/10) | CO | - | 173.2/171.1 | | ζ | 7.18 | 126.1 |
| | N | 7.59/7.64 | - | Arg (13) | CO | - | 172.1 |
| | α | 4.19/4.31 | 57.2/56.4 | | N | 8.20 | - |
| | β | 1.97/1.91 | 30.6/30.5 | | α | 4.19 | 51.2 |
| | γ | 0.82, 0.86/ 0.84, 0.87 | 17.7/19.0 | | β | 1.60, 1.75 | 27.9 |
| Gly (5) | CO | - | n.d. | | γ | 1.49 | 24.7 |
| | N | 8.08 | - | | δ | 3.09 | 40.0 |
| | α | 3.69, 3.80 | 42.3 | | Nε | 7.67 | - |

[a]800 MHz; [b]Determined by HSQC and HMBC at 800 MHz; n.d., not detected.

**Table S4.** Primers used for generating *ΔmpcB::apr*$^{R}$ in *S. ghanaensis*. Sequences are shown in the 5’-3’ direction. Recognition sites of restriction enzymes are underlined.

| Primer | Enzyme | Sequence |
|---|---|---|
| Sg_ΔB12rSAM_L_for | SpeI | ATATATACTAGTAGCGAGGGGTTCAGCGTGCACC |
| Sg_ΔB12rSAM_L_rev | EcoRI | ATATATGAATTCAGAGCCAGCGACGGGAGATCGATCG |
| Sg_ΔB12rSAM_R_for | HindIII | ATATATAAGCTTAGCGGGCAGTTCGTGCAGATCGCG |
| Sg_ΔB12rSAM_R_for | KpnI | ATATATGGTACCATGGCCTGGGTGCGGGACTTGATCG |
| Sg_ΔB12rSAM_check_for | - | TGACGATTTCCCGGCCGACGC |
| Sg_ΔB12rSAM_check_rev | - | TGTGCCGCATGTGGTCGATCCGC |

**Table S5.** Primers used for cloning genes from the *S. clavuligerus* genome for heterologous expression in *E. coli*. Sequences are shown in the 5'-3' direction. Recognition sites of restriction enzymes are underlined. Ribosomal binding site is shown in bold.

| Primer | Enzyme | Sequence |
|---|---|---|
| MpcA_for | BamHI | ATATATGGATCCATGCAGAAGGACGTCATTCACAACG |
| MpcA_rev | HindIII | ATATATAAGCTTTCAGTCCTCGTCCGCGTCCTC |
| MpcB_for | NdeI | ATATATCATATGATGCGTGTTCTGCTGGTCAATATGCC |
| MpcB_rev | KpnI | ATATATGGTACCTCATACGTGAACGGGTTCGGCCG |
| MpcC_for | XhoI | ATCTCGAG**AATAAGGAGGTATACA**<br>ATGAGCACCGCCCTGAGTCCG |
| MpcC_rev | XhoI | ATATACTCGAGTCAGATCGTCATCGACTCCTGGAGCC |
| MpcC_for_GA | - | GAACCCGTTCACGTATGAGGTACCC<br>**AATAAGGAGGTATACA**ATGAGCACCGCCCTG |
| MpcC_rev_GA | - | GCAGCGGTTTCTTTACCAGACTCGA<br>TCAGATCGTCATCGACTCCTGGAGCC |
| MpcC_for_NdeI | NdeI | ATATATCATATGATGAGCACCGCCCTGAGTCCG |
| MpcA_rev_X | XhoI | ATATATCTCGAGTCAGTCCTCGTCCGCGTCCTC |
| MpcP_for | NdeI | AAATTTCATATGATGAGCTATCAGGTCTATCGCGCGG |
| MpcP_rev | XhoI | ATATATCTCGAGTCACCCGCGAGCGGGTCGTTG |

**Table S6.** Tandem MS fragments of octa-methylated $MpcA_{1-13}$ peptide from heterologous expression of MBP-MpcA and MpcBC in *E. coli*.

| Ion | #Methylations | m/z obs. | m/z calc. | Δ (ppm) |
|---|---|---|---|---|
| $b1^+$ | 0 | 129.1040 | 129.1023 | 13.2 |
| $b2^+$ | 2 | 254.1909 | 254.1864 | 17.7 |
| $b4^+$ | 3 | 454.3092 | 454.3024 | 15.0 |
| $b6^+$ | 4 | 638.4277 | 638.4236 | 6.4 |
| $y1^+$ | 0 | 175.1223 | 175.1190 | 18.8 |
| $y2^+$ | 0 | 322.1930 | 322.1874 | 17.4 |
| $y3^+$ | 2 | 447.2802 | 447.2715 | 19.5 |
| $y9^+$ | 5 | 1073.6866 | 1073.6718 | 13.8 |
| $y11^+$ | 6 | 1273.8043 | 1273.7879 | 12.9 |

**Table S7.** Distribution of 'long' precursor peptides with acidic followers and 'short' precursor peptides without acidic followers in Actinobacteria.

| Strains | # 'long' precursors | # 'short' precursors |
|---|---|---|
| *Streptomyces* | 204 | 2 |
| Other Actinobacteria genera | 2 | 20 |

**Figure S1.** Results of HHpred alignment of MpcC (WP_003958122.1) with MpcB (WP_003958123.1), indicating structural similarities between these 2 proteins.

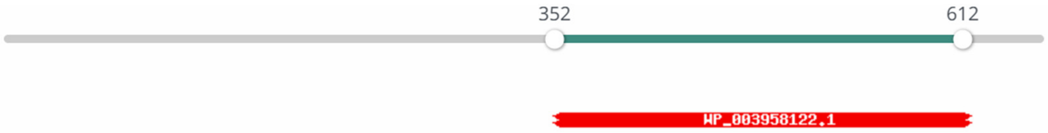


### Hitlist

Show 25 Entries Search:

| Nr | Hit | Name | Probability | E-value | Score | SS | Aligned cols | Target Length |
|---|---|---|---|---|---|---|---|---|
| 1 | WP_003958122.1 | DUF5825 family protein [Streptomyces clavuligerus] | 99.78 | 1.7e-22 | 168.01 | 18.9 | 173 | 221 |

**Figure S2.** PCR validation of $P_{ermE}$* insertion into *S. clavuligerus* using primers *ScvIns_check_for* and *ScvIns_check_rev*. Expected PCR product lengths for WT and engineered strains are 588 bp and 391 bp, respectively.

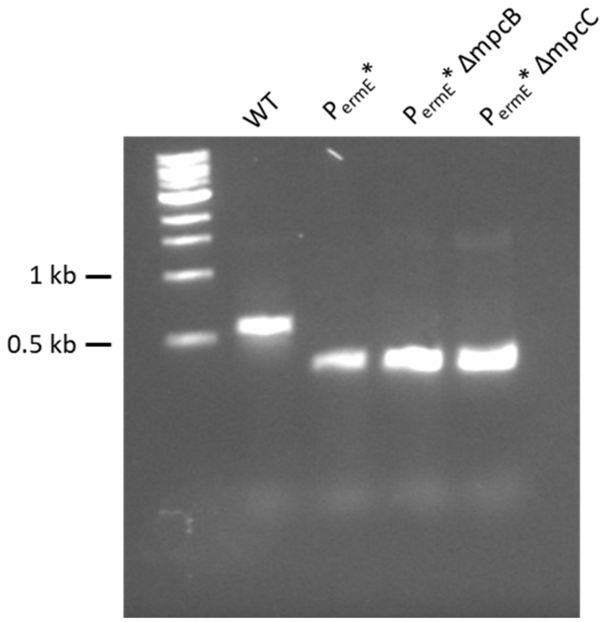


**Figure S3.** PCR verification of *mpcB* (left) and *mpcC* (right) knockouts of engineered *S. clavuligerus* $P_{mpc}$::$P_{ermE}$* using primers *MpcB_for* and *MpcB_rev*, and *MpcC_for* and *MpcC_rev*, respectively. Expected band lengths are marked.

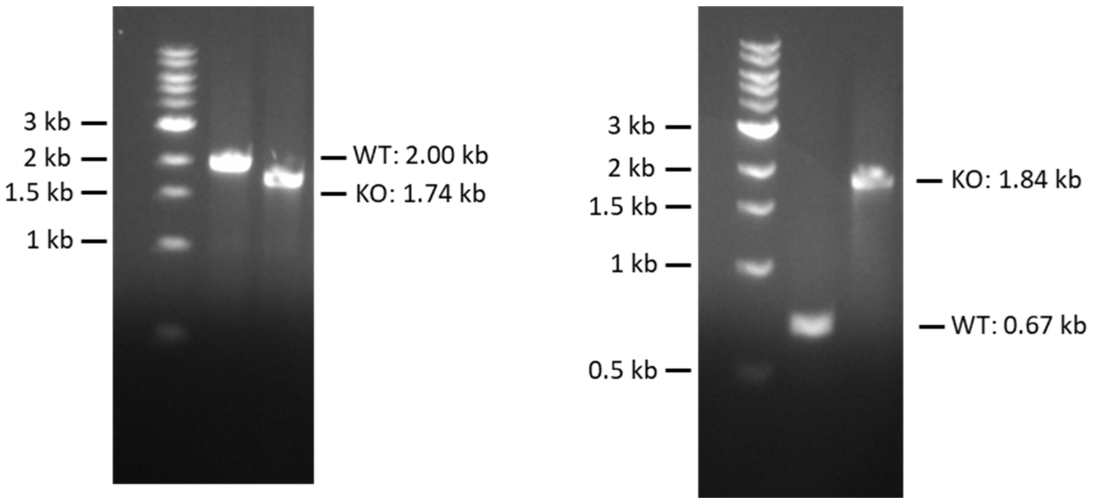

**Figure S4.** 800 MHz NMR spectra of clavusporins (with variant B as the major component) in DMSO-$d_6$. $^{1}H$, COSY, TOCSY, HSQC, and HMBC spectra are shown from top to bottom.

$^{1}H$

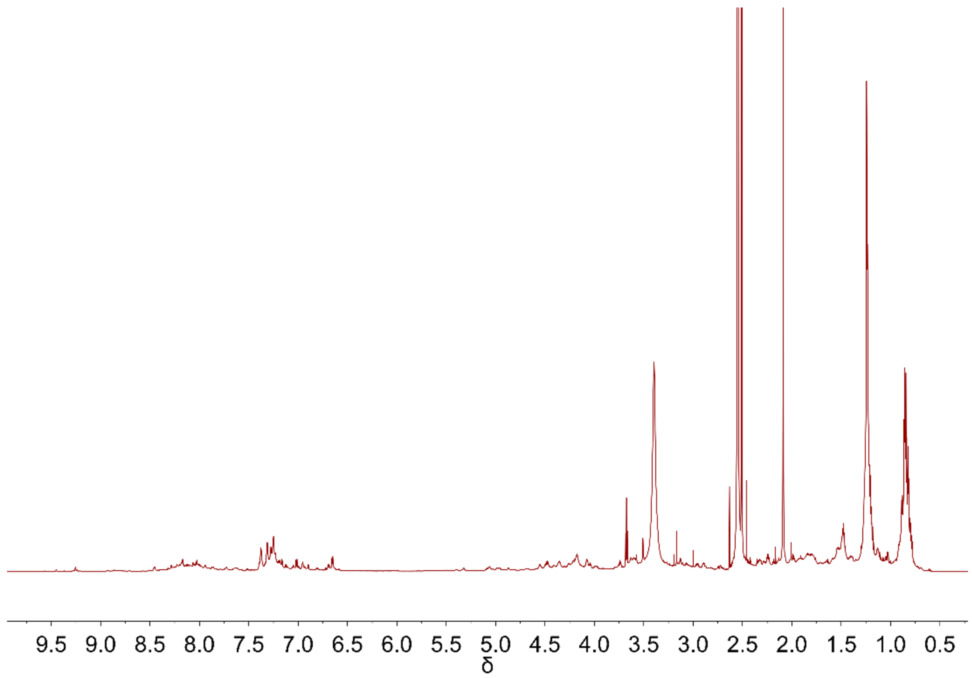


COSY

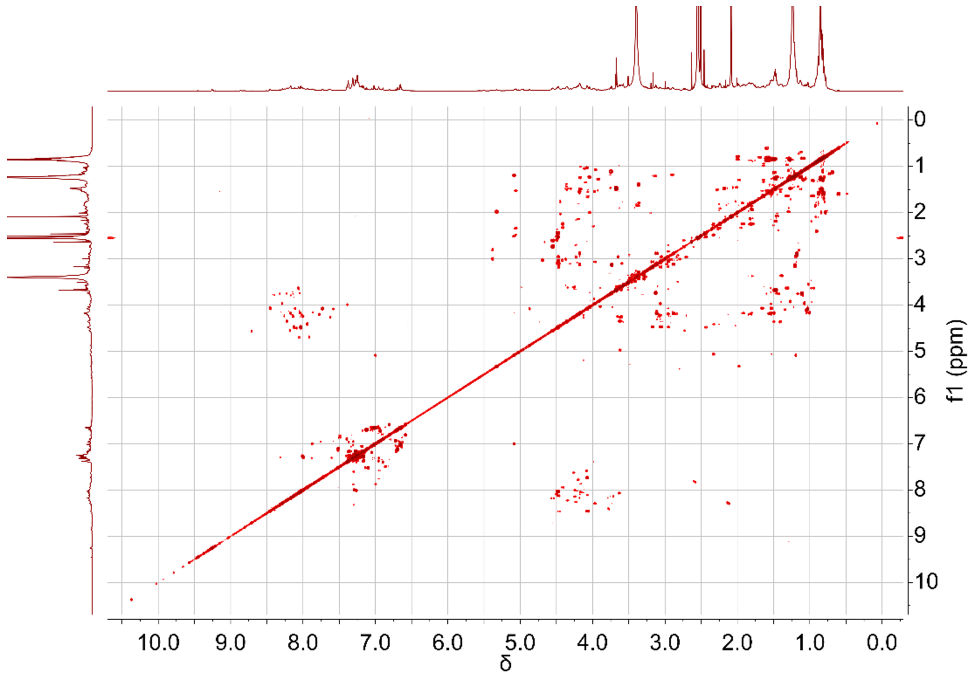

TOCSY

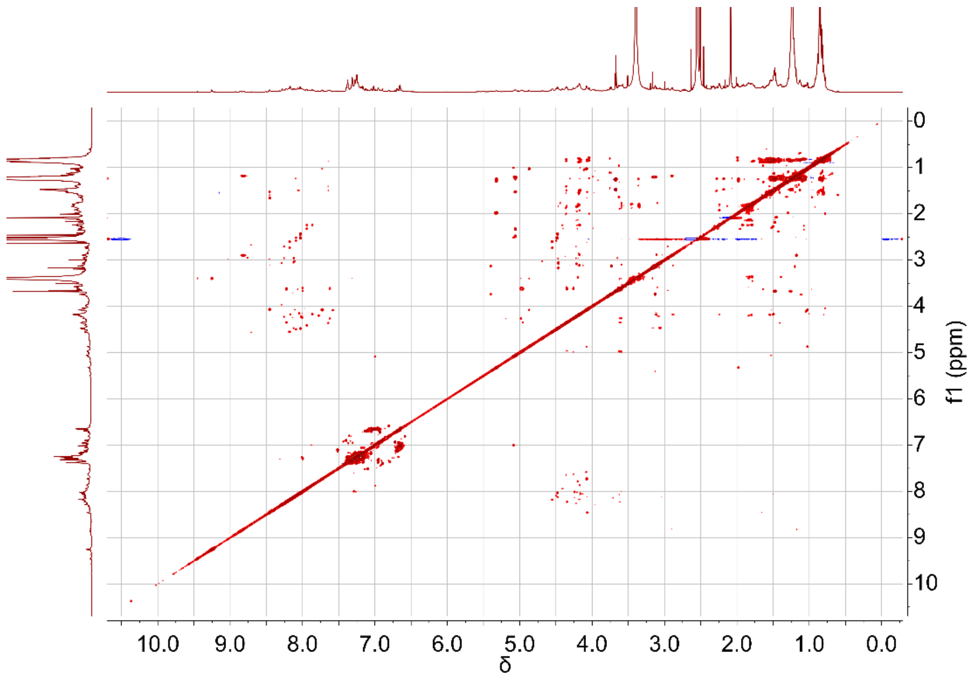

0
1
2
3
4
5
6
7
8
9
10
f1 (ppm)
10.0
9.0
8.0
7.0
6.0
5.0
4.0
3.0
2.0
1.0
0.0
δ


HSQC

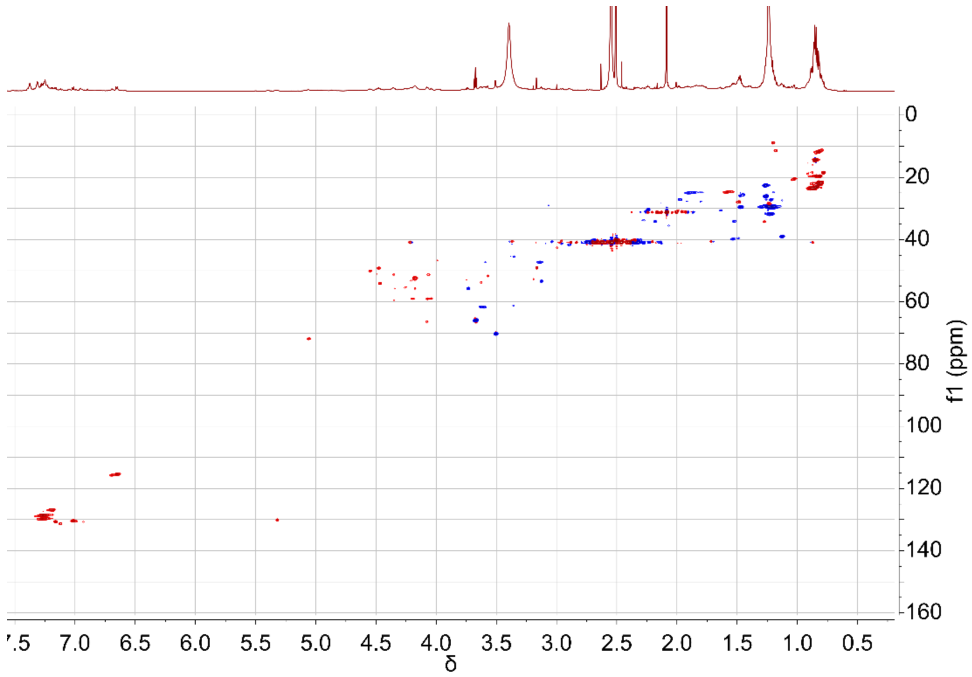

0
20
40
60
80
100
120
140
160
f1 (ppm)
7.0
6.5
6.0
5.5
5.0
4.5
4.0
3.5
3.0
2.5
2.0
1.5
1.0
0.5
δ

HMBC

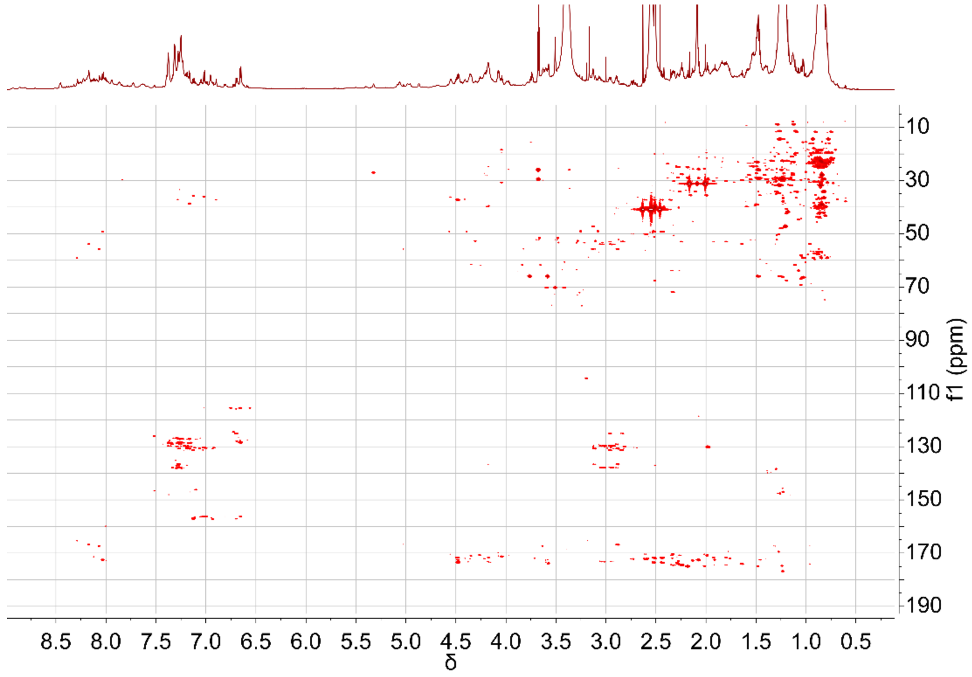
10
30
50
70
90
110
130
150
170
190
f1 (ppm)
8.5
8.0
7.5
7.0
6.5
6.0
5.5
5.0
4.5
4.0
3.5
3.0
2.5
2.0
1.5
1.0
0.5
δ

**Figure S5.** 800 MHz NMR spectra of unmodified $MpcA_{1-13}$ (core peptide) in DMSO-$d_6$. $^1H$, COSY, TOCSY, HSQC, and HMBC spectra are shown from top to bottom. The key impurity is the $CF_3COOH$ residual.

<u>$^1H$</u>

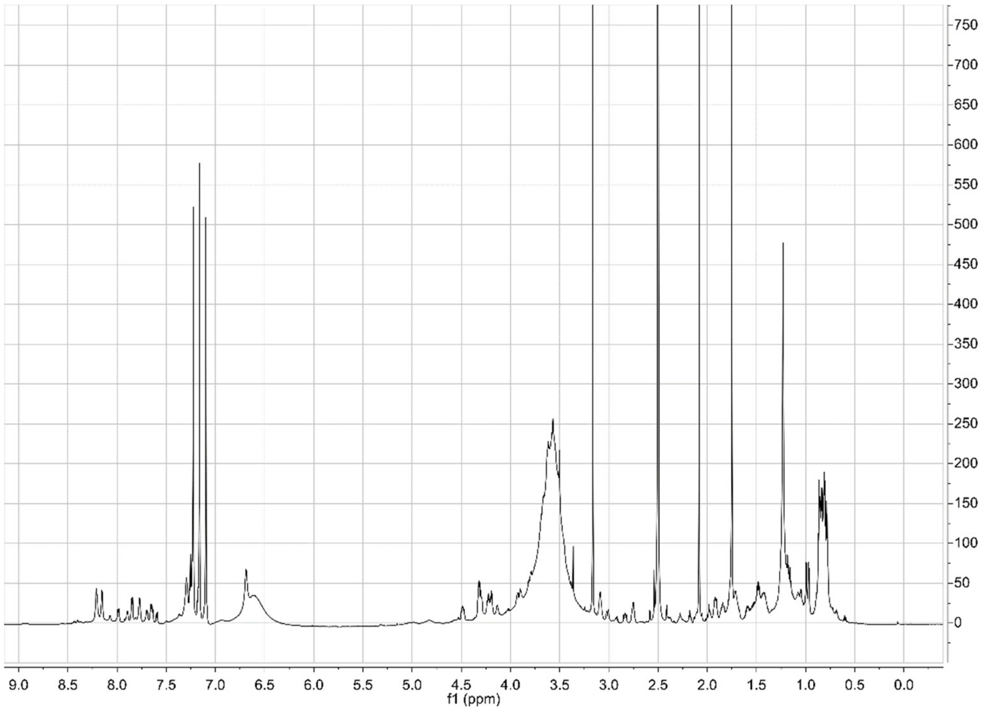

### COSY

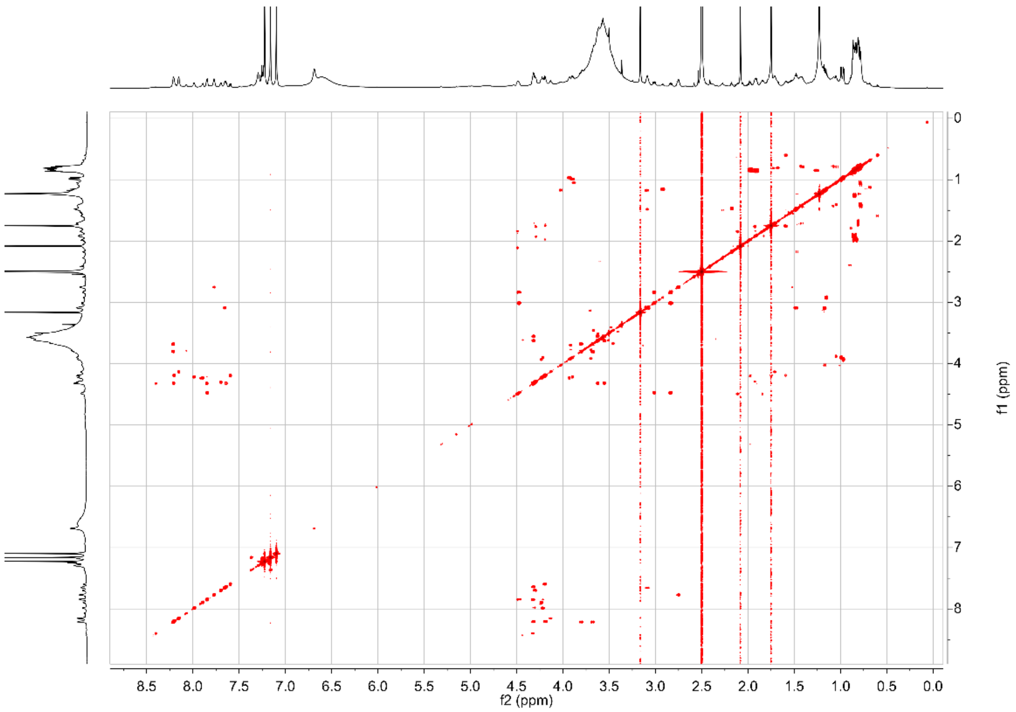


### TOCSY

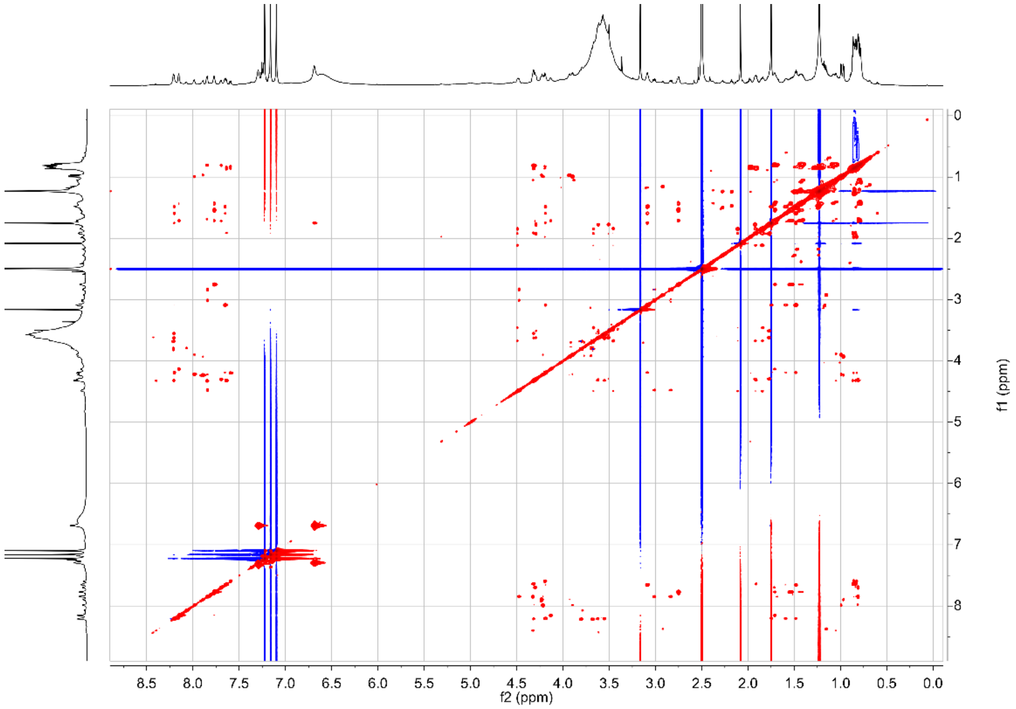

### HSQC

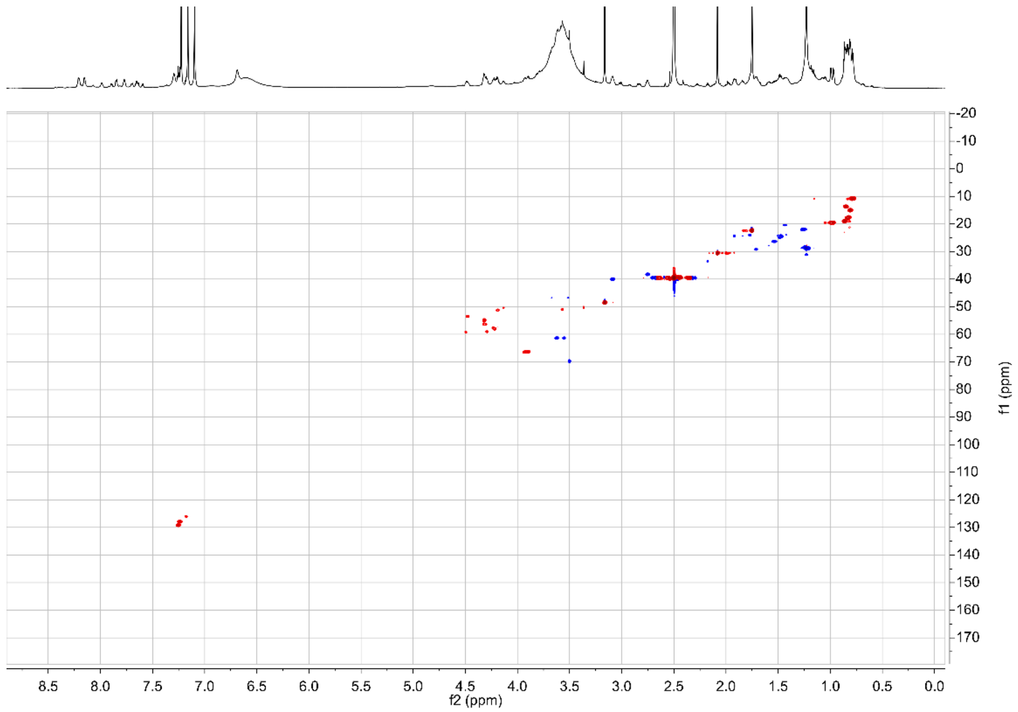

### HMBC

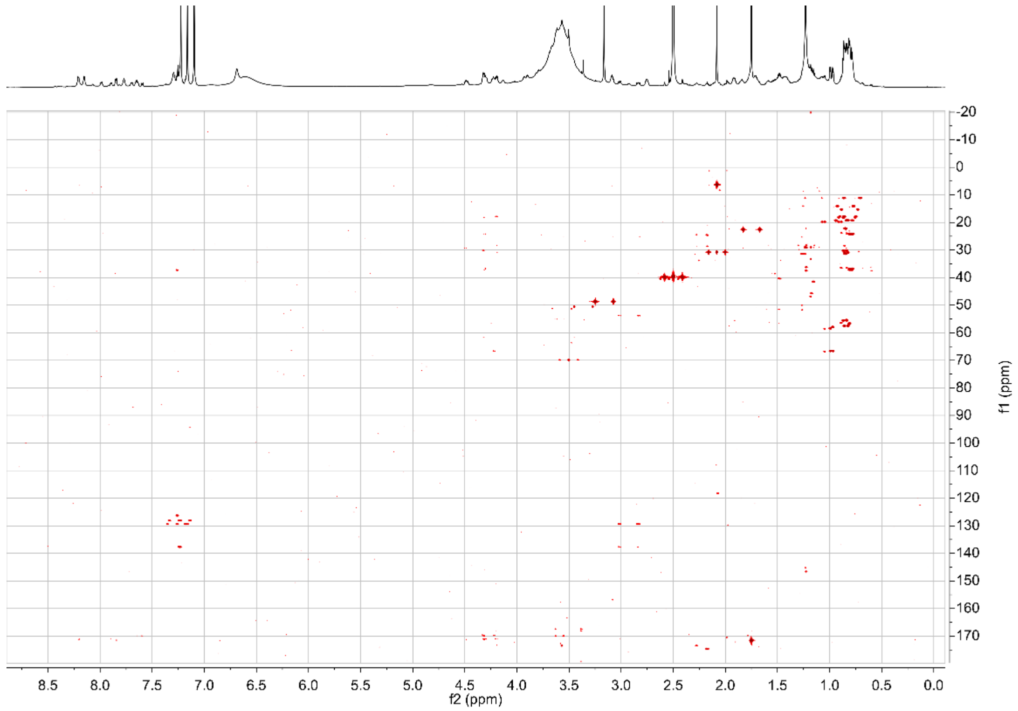

**Figure S6.** PCR verification of *mpcB* (left) and *mpcC* (right) knockouts of *S. clavuligerus* using primers *MpcB_for* and *MpcB_rev*, and *MpcC_for* and *MpcC_rev*, respectively (Table S5). Expected band lengths are marked.

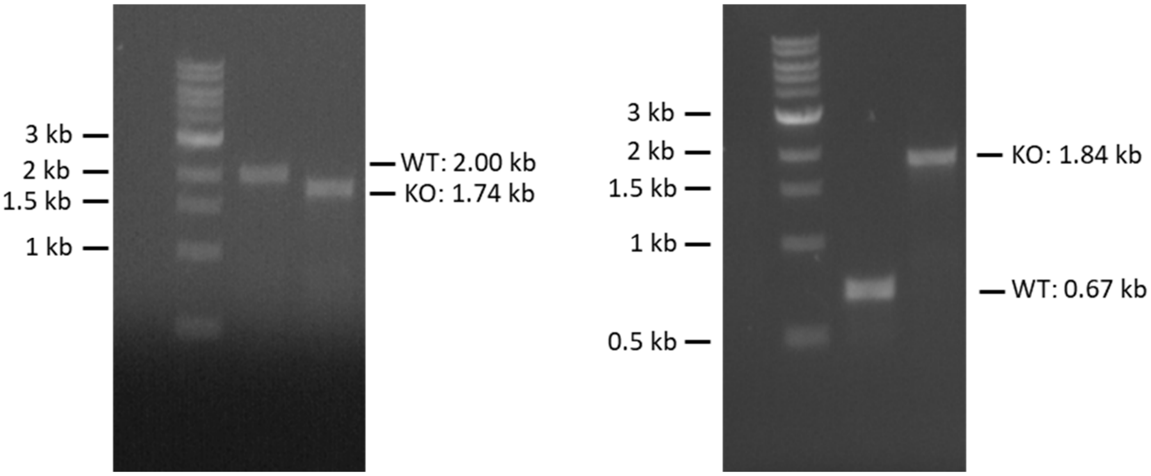


**Figure S7.** Compound complementation of various amounts to clavusporin-deficient *S. clavuligerus* mutants, showing concentration-dependent promotion of aerial hyphae growth.

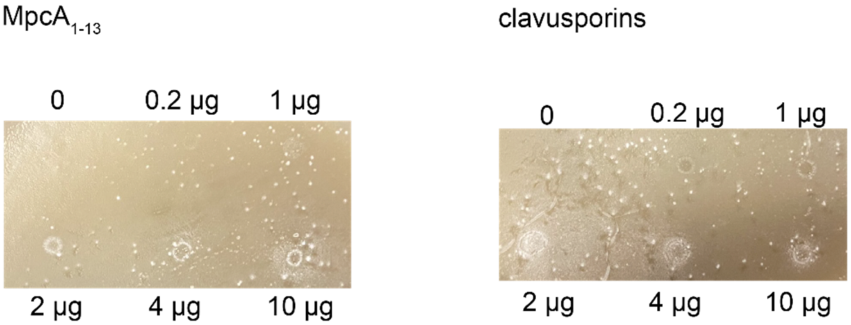

**Figure S8.** PCR verification of *mpgB* knockout in WT *S. ghanaensis* using primers *Sg_ΔB12rSAM_check_for* and *Sg_ΔB12rSAM_check_rev* (Table S6). Expected band lengths are marked.

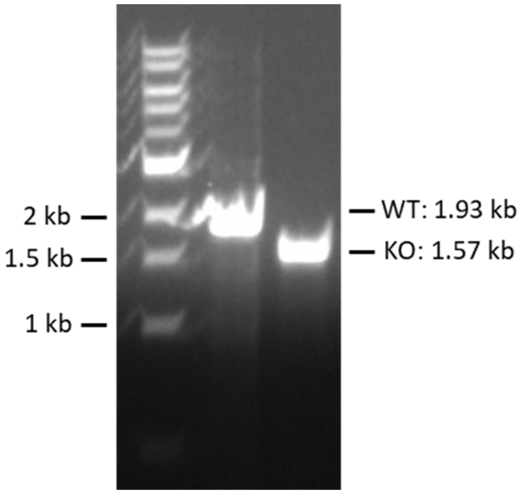
